# A latent hydrazone switch turns SP2509 into an optical timer of cell fate

**DOI:** 10.64898/2026.08.04.742687

**Authors:** Chuan-Shuo Wu, Li Liu, Liang Cheng

## Abstract

Epigenetic inhibitors can reprogram cell states, but their activity is usually governed by fixed dose-response relationships rather than by user-defined temporal commands. Here we show that **SP2509**, an established reversible LSD1/KDM1A inhibitor, contains a latent *ortho*-hydroxy acylhydrazone switch that converts this known scaffold into an optical timer for chromatin-dependent cell fate. Irradiation at 430 nm enriches a less active *cis*-associated state under near-physiological conditions, whereas dark relaxation regenerates the more active *trans*-associated state with a half-life of 6.5 hours. This reversible conformational change reduces recombinant LSD1 inhibition by approximately 7-fold and is transmitted to H3K9me2 accumulation, proliferation, senescence and apoptosis. Reciprocal light-dark switching experiments reveal a reversible window for senescence-associated arrest in MGC-803 cells and tune population-level entry into apoptosis in OCI-AML3 cells. RNA-seq endpoint comparisons further distinguish photostate-associated transcriptional states, with dark *trans*-**SP2509** producing stronger stress-survival and apoptosis-linked programs than cells maintained under *cis*-**SP2509** conditions. These findings establish latent hydrazone photoisomerization as a compact strategy for converting a static epigenetic inhibitor into a kinetic regulator of chromatin-dependent cell-fate transitions.

---

Chromatin-modifying enzymes convert transient biochemical inputs into transcriptional states that can persist beyond the initiating perturbation. Lysine-specific demethylase 1 (LSD1/KDM1A) exemplifies this principle because it functions in distinct cofactor-dependent complexes and contributes to transcriptional repression, gene activation, lineage control and tumor-cell plasticity. Its originally described activity removes mono- and dimethyl marks from H3K4 in repressive complexes, whereas later studies linked LSD1-containing assemblies to H3K9 demethylation in nuclear-receptor contexts. This context dependence makes LSD1 an attractive chemical target,^1,2^ but it also means that the biological consequence of inhibition is likely to depend not only on inhibitor potency, but also on when and for how long the enzyme is perturbed during chromatin-state evolution.^3–6^ The chemical problem is therefore not simply to inhibit LSD1, but to control the timing of LSD1 inhibition during an evolving cellular trajectory.^7,8^ Irreversible catalytic inhibitors and reversible scaffolding inhibitors have provided powerful ways to perturb LSD1-dependent transcription, differentiation and tumor-cell survival.^9,10^ However, conventional dosing fixes inhibitor exposure at the beginning of an experiment and cannot subsequently alter target engagement within the same culture. This limitation is especially important for chromatin enzymes, where transient differences in perturbation history may determine whether cells compensate, enter reversible arrest or cross an irreversible commitment threshold.^11–13^

Photo-pharmacology offers a route to temporal control by using light to interconvert ligands between states with different biological activities.^14–20^ Most established systems install a dedicated photochrome, such as azobenzene, diazocine or hemithioindigo, into a lead molecule.^18–24^ This strategy can be powerful, but adding an external switching module may alter potency, permeability, selectivity or metabolic behavior. An alternative strategy is to search existing bioactive scaffolds for latent photochemical elements that have been treated only as pharmacophoric features. Such an approach would preserve the drug-like architecture of the parent compound while adding a temporal command layer.^25–28^

The *ortho*-hydroxy acylhydrazone in **SP2509** places this strategy in a chemically tractable framework. Acylhydrazones can undergo C=N isomerization, and *ortho*-hydroxy substitution can couple hydrazone geometry to proton transfer, intramolecular hydrogen bonding and thermal relaxation. Previous studies of bioactive acylhydrazones have shown that light or thiols can alter isomer distributions and that different isomers can display distinct biological activities.^29–32^ We therefore asked whether this latent dynamic element in **SP2509** could be uncovered as a functional photochemical switch for timing chromatin-enzyme inhibition.^33,34^ Here, *trans*- and *cis*-refer to the configuration about the hydrazone C=N bond. We show that the *trans*-geometry of **SP2509** is the more active LSD1-inhibitory state, whereas 430 nm light enriches a less active *cis*-ensemble that thermally relaxes in the dark. This molecular switch modulates recombinant enzyme inhibition, H3K9me2 accumulation, cell growth, senescence and apoptosis. RNA-seq endpoint comparisons identify photostate-associated transcriptional states, and reciprocal switching experiments show that photostate history controls the rate at which cells enter senescence or apoptosis. Together, these results define **SP2509** as a compact optical timer for interrogating how the timing of LSD1 inhibition shapes tumor-cell fate.

## RESULTS

### A latent hydrazone switch is revealed in SP2509

The chemical premise is illustrated by a general hydrazone model (Fig. 1a). The *trans*-geometry separates the larger substituents (R^L^) across the C=N bond and generally adopts a more extended arrangement, whereas irradiation at one wavelength (λ_1_) can populate a more compact *cis*-geometry. A second wavelength (λ_2_) or thermal relaxation (Δ) can drive the reverse *cis*→*trans* process, creating a reversible optical or photothermal cycle rather than a one-way structural change.

**Fig. 1.**
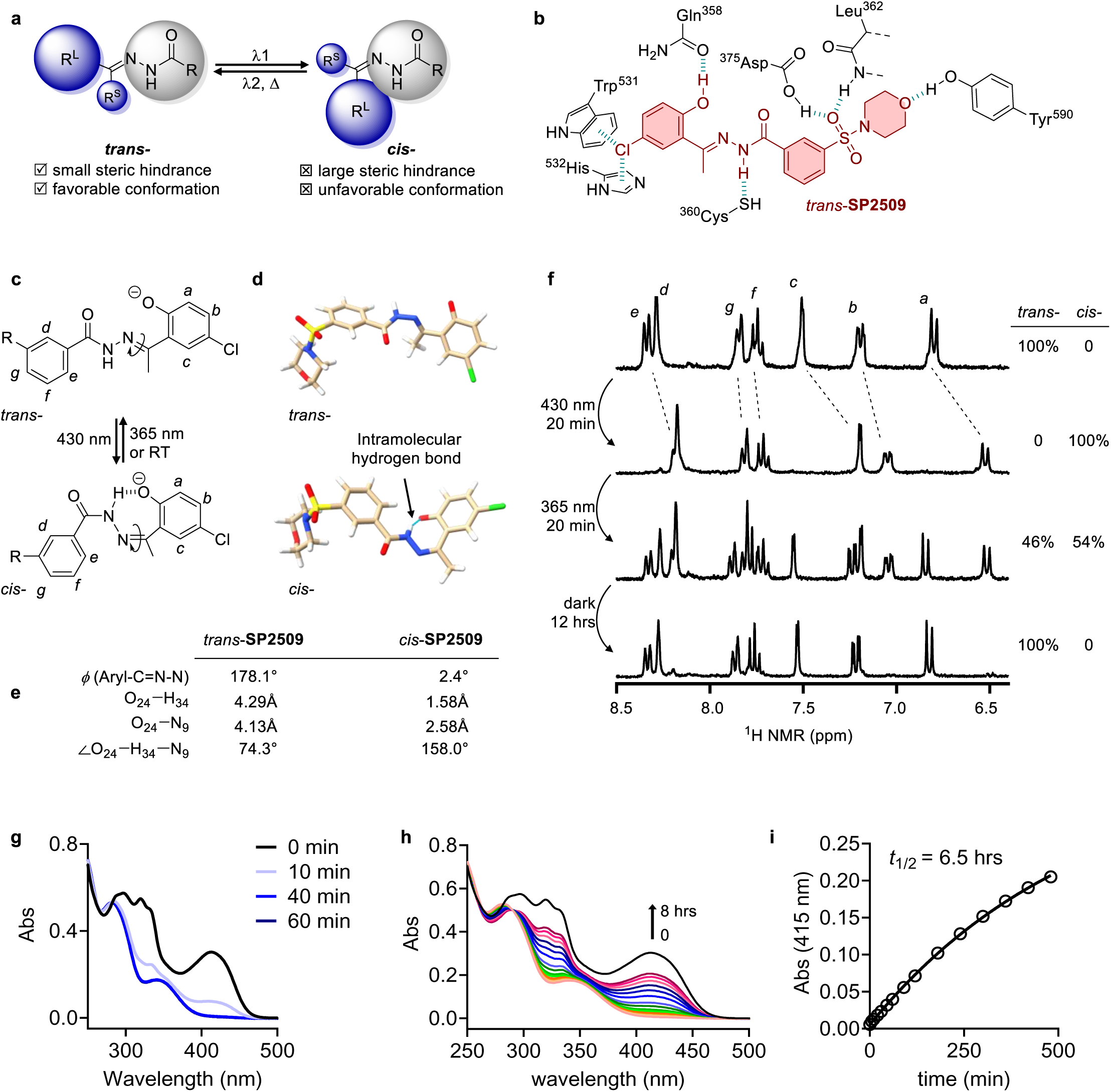
A latent *ortho*-hydroxy acylhydrazone switch in SP2509 enables reversible optical isomerization. **a**, General schematic of hydrazone *cis*/*trans*-photoisomerization. The *trans*-configuration separates the larger substituents (R^L^) and generally adopts a more extended arrangement, whereas irradiation can populate a compact *cis*-configuration; a second wavelength or heat drives the reverse process. **b**, Proposed binding model of *trans*-**SP2509** in the LSD1 pocket, showing an extended geometry that positions the chloroaryl, phenolic, hydrazone, sulfonamide and morpholine-containing regions near polar and aromatic residues that can support a distributed interaction network. **c**, Proposed photoisomerization scheme of deprotonated **SP2509**, showing 430 nm-induced *trans*-to-*cis* conversion and reverse *cis*-to-*trans* recovery by 365 nm irradiation or thermal relaxation at room temperature. **d**, DFT-optimized geometries of the deprotonated phenolate forms of *trans*-and *cis*-**SP2509**. Both structures were confirmed as local minima by harmonic frequency analysis. The *trans*-isomer adopts an open geometry, whereas the *cis*-isomer folds into a pseudocyclic conformation stabilized by an intramolecular N–H···O(phenolate) hydrogen bond. **e**, Comparison of selected structural parameters. The absolute N–N=C–Ar dihedral changes from 178.1°in *trans*-**SP2509** to 2.4°in *cis*-**SP2509**. The corresponding O···H and O···N distances change from 4.29 and 4.13 Å to 1.58 and 2.58 Å, respectively. Calculations were performed at the ωB97X-D/6-31G(d) level using IEFPCM solvation in DMSO. **f**, ^1^H NMR analysis of **SP2509** photoisomerization in DMSO-*d*_6_ after triethylamine addition. 430 nm irradiation for 20 minutes converted the *trans*-dominant spectrum to a *cis*-enriched spectrum, 365 nm irradiation partially restored the *trans*-state, and dark incubation for 12 hours regenerated the *trans*-dominant state. **g**, UV-vis absorption spectra of **SP2509** in aqueous buffer under 430 nm irradiation, showing depletion of the *trans*-associated absorption band around 413 nm and emergence of a band around 341 nm. **h**, UV-vis spectra monitoring thermal relaxation of *cis*-enriched **SP2509** in the dark. **i**, Thermal relaxation kinetics monitored by recovery of absorbance at 415 nm in phosphate-buffered saline at room temperature, giving a half-life of approximately 6.5 hours.

We synthesized the *trans*-isomer of **SP2509** by the established route and confirmed its structure by nuclear magnetic resonance (NMR) spectroscopy and high-resolution mass spectrometry (Fig. 1b and Supplementary Fig. S1). **SP2509** contains an *ortho*-hydroxy acylhydrazone that matches this switching logic, with the hydrazone C=N bond and adjacent phenol positioned to support proton-coupled *trans*/*cis*-photoisomerization. In the proposed LSD1 binding model, *trans*-**SP2509** adopts an extended geometry that spans the pocket and positions its chloroaryl, phenolic, hydrazone, sulfonamide and morpholine-containing regions within a distributed network of polar and aromatic residues (Fig. 1b,c).^12,35,36^

We next asked whether the proposed binding difference could arise from an intrinsic reorganization of the **SP2509** pharmacophore. DFT calculations on the deprotonated phenolate forms of *trans*- and *cis*-**SP2509** identified distinct local-minimum geometries, with no imaginary frequencies (Fig. 1d,e and Supplementary Fig. S2, Supplementary Tables S1 and S2). The absolute N–N=C–Ar dihedral changed from 178.1°in the *trans*-isomer to 2.4°in the *cis*-isomer. Whereas *trans*-**SP2509** retained an open geometry, *cis*-**SP2509** folded into a pseudocyclic conformation stabilized by an intramolecular N–H···O(phenolate) hydrogen bond (O···H, 1.58 Å; O···N, 2.58 Å; N–H···O, 158.0°). This isomer-dependent folding provides a plausible structural rationale for the reduced LSD1 inhibitory activity of the *cis*-enriched state.

We next tested whether this latent hydrazone switch could be activated by light using ¹H NMR spectroscopy (Fig. 1f and Supplementary Fig. S3). Addition of triethylamine caused the exchangeable phenolic and hydrazide NH resonances to disappear, consistent with formation of a base-responsive deprotonated ensemble. Irradiation at 430 nm for 20 minutes drove conversion from a *trans*-dominant spectrum (*trans*:*cis* = 100:0) to a *cis*-enriched spectrum (*trans*:*cis* = 0:100). Subsequent irradiation at 365 nm partially restored the *trans* state (*trans*:*cis* = 46:54), and dark incubation for 12 hours returned the sample to a *trans*-dominant spectrum (*trans*:*cis* = 100:0) (Fig. 1f). These data establish that deprotonated **SP2509** undergoes reversible, visible-light-responsive hydrazone isomerization.

We then asked whether this behaviour persists under conditions compatible with biological experiments. In phosphate-buffered saline containing 10% DMSO, 50 µM of **SP2509** displayed a yellow color and a pronounced absorption band at approximately 413 nm (Supplementary Fig. S4). Irradiation at 430 nm (2 mW·cm^−2^) progressively decreased this band and generated a new band at approximately 341 nm, reaching an apparent photostationary state after approximately 40 minutes (Fig. 1g). In the dark, the spectrum recovered toward the initial *trans*-associated profile, and fitting the recovery of absorbance at 415 nm gave a thermal half-life of approximately 6.5 hours at room temperature (Fig. 1h,i). Thus, the embedded hydrazone switch can be optically attenuated under aqueous conditions and thermally restored on a timescale suitable for cell-fate experiments. Together, these structural, spectroscopic and computational analyses establish that **SP2509** contains a latent hydrazone photoswitch that reversibly reorganizes its pharmacophore without introducing an external photoswitch scaffold.

### Photoisomerization gates LSD1 inhibition and chromatin output

We next asked whether the hydrazone geometry changes target inhibition. In a recombinant LSD1 assay, *trans*-**SP2509** inhibited the enzyme with low-nanomolar potency, whereas a 430 nm pre-irradiated, *cis*-enriched sample was substantially less active (Fig. 2a,b and Supplementary Fig. S5). Across independent measurements, the *trans*- and *cis*-enriched states gave IC_50_ values of 24.9 nM and 177.9 nM, respectively, corresponding to an approximately 7-fold optical difference in enzymatic potency. This result is consistent with a model in which the extended *trans*-geometry better complements the proposed LSD1-binding pocket than the compact, intramolecularly hydrogen-bonded *cis*-geometry (Fig. 1b,d,e).

**Fig. 2.**
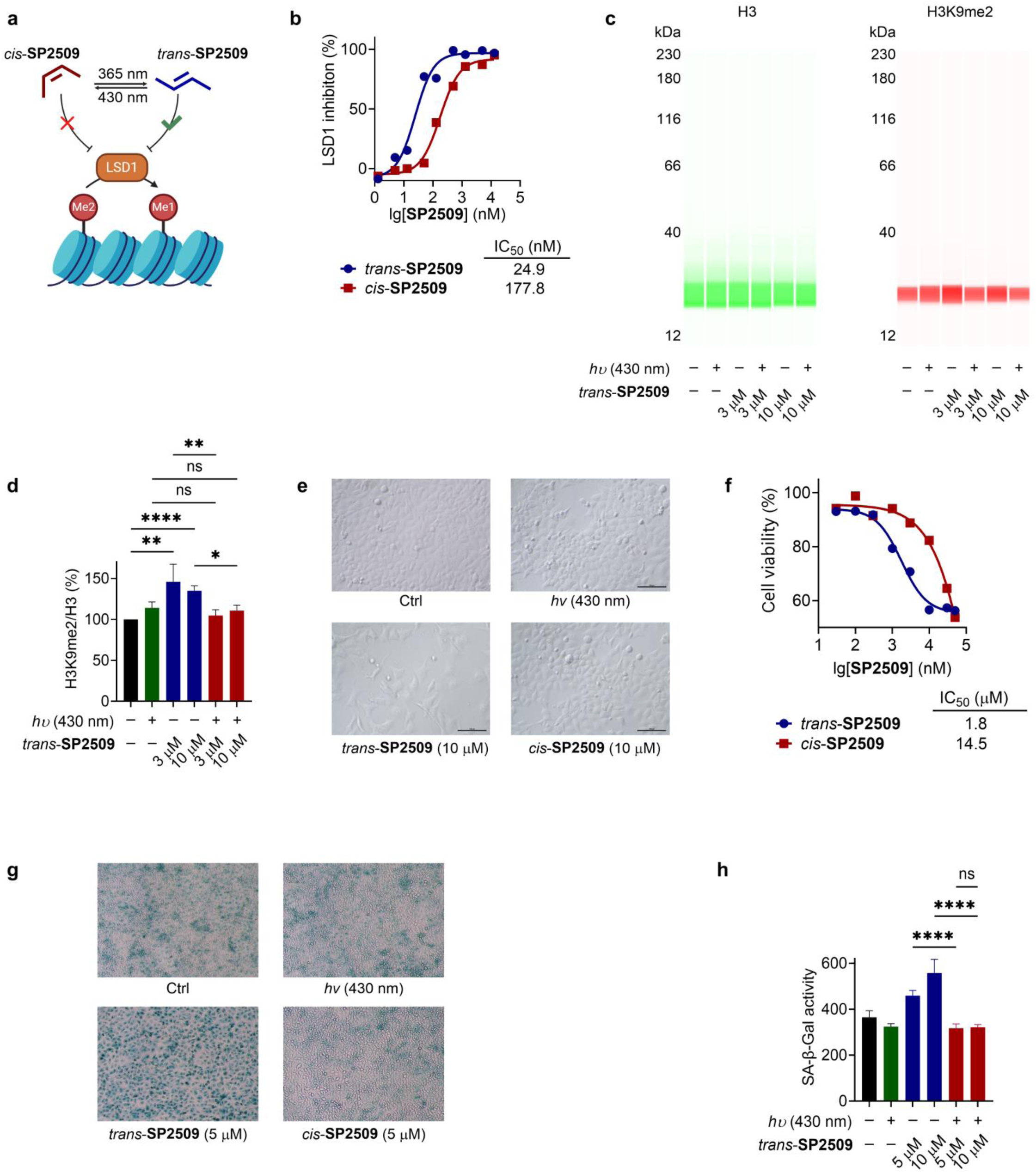
Photoconversion of SP2509 gates LSD1 inhibition, chromatin output and senescence-associated phenotypes. **a**, Schematic showing photostate-dependent regulation of LSD1 by **SP2509**. *Trans*-**SP2509** functions as the more active LSD1-inhibitory state, whereas 430 nm illumination enriches a less active *cis*-state. **b**, Recombinant LSD1 inhibition curves for *trans*- and *cis*-enriched **SP2509**, with IC_50_ values of 24.9 nM and 177.8 nM, respectively. **c**, Representative Simple Western analysis of H3K9me2 and total H3 in HeLa cells treated with **SP2509** under dark or 430 nm illuminated conditions. **d**, Quantification of H3K9me2 normalized to total H3, showing a 35–46% increase after *trans*-**SP2509** treatment and a weaker response in cells maintained under *cis*-enriched **SP2509** conditions at matched concentrations. **e**, Representative bright-field images of MGC-803 cells treated with vehicle, 430 nm illumination alone, *trans*-**SP2509** or *cis*-enriched **SP2509**. **f**, Cell-viability analysis showing stronger growth inhibition by *trans*-**SP2509** than by *cis*-enriched **SP2509** after 48 hours. **g**, Representative senescence-associated β-galactosidase staining images of MGC-803 cells under the indicated conditions. **h**, Quantification of senescence-associated β-galactosidase activity, showing stronger senescence-associated activity in cells treated with dark *trans*-**SP2509** than in cells maintained under *cis*-enriched **SP2509** conditions. Data are presented as mean ±s.e.m. from independent biological replicates. Statistical significance was determined using an unpaired two-tailed Student’s *t*-test. Statistical significance is indicated as follows: ns, not significant; \**P* < 0.05; \*\**P* < 0.01; \*\*\**P* < 0.001; \*\*\*\**P* < 0.0001.

We then tested whether this biochemical difference reaches chromatin. HeLa cells were treated with **SP2509** under dark (*trans*-) or 430 nm illuminated (*cis*-enriched) conditions, and H3K9me2 was quantified by Simple Western using total H3 as a loading control. *Trans*-**SP2509** increased H3K9me2 by 35–46% relative to vehicle and light-only controls, consistent with cellular LSD1 inhibition in this context (Fig. 2c,d). At matched compound concentrations and endpoint conditions, cells maintained under *cis*-enriched **SP2509** conditions showed a weaker H3K9me2 response of approximately 5–10%, whereas low-intensity intermittent 430 nm illumination alone produced only a minor change (Fig. 2c,d). Thus, the photochemical state of **SP2509** is transmitted from molecular geometry to enzyme output and then to a chromatin mark.

The same photostate dependence was evident at the level of cell growth and senescence-associated phenotype (Fig. 2e-h). In MGC-803 gastric cancer cells, 10 µM of *trans*-**SP2509** reduced cell number after 48 hours, whereas *cis*-enriched **SP2509** produced weaker growth inhibition under matched illumination conditions (Fig. 2e). Cell-viability curves revealed an 8-fold shift in growth-inhibitory potency, with IC_50_ values of 1.8 µM and 14.5 µM for *trans*- and *cis*-enriched **SP2509**, respectively (Fig. 2f). Senescence-associated (SA) β-galactosidase staining and quantitative senescence-associated β-galactosidase activity measurements further showed that *trans*-**SP2509** induced a senescence-associated state (26–53%), whereas cells maintained under *cis*-enriched **SP2509** conditions showed markedly reduced activity (-12–-13%) (Fig. 2g,h). Hydrazone geometry is therefore converted into a quantitative difference in LSD1 inhibition that is large enough to reach chromatin and reshape tumor-cell phenotype.

### Optical timing defines a reversible senescence window

A useful optical probe should do more than compare two static photostates. It should allow target inhibition to be moved relative to a biological trajectory.^37,38^ We therefore performed reciprocal switching experiments in MGC-803 cells (Fig. 3). In the *trans*-to-*cis* protocol, cells were exposed to *trans*-**SP2509** for 24 hours to inhibit LSD1 and initiate a senescence-associated program (Fig. 3a and Supplementary Fig. S7, S8, S11). At this stage, cells treated with 2 or 5 µM of *trans*-**SP2509** displayed flattened morphology, enlarged nuclei, increased senescence-associated β-galactosidase activity and S-phase accumulation (Fig. 3b, 3c).^39^ Cultures were then either maintained in the dark for another 42 hours or transferred to 430 nm illumination to initiate the *trans*→*cis* transition. Continuous dark treatment preserved the senescence-associated state after 66 hours (Fig. 3d, 3e), whereas the light-shifted condition reduced senescence-associated β-galactosidase activity and partially restored the cell-cycle profile (Fig. 3f, 3g). Optical attenuation after senescence initiation therefore exposes a reversible decision window in which LSD1 inhibition has initiated a senescence program (Fig. 3h, 3i), but the chromatin and cell-cycle states remain sufficiently plastic to be redirected by changing the photostate of **SP2509**.

**Fig. 3.**
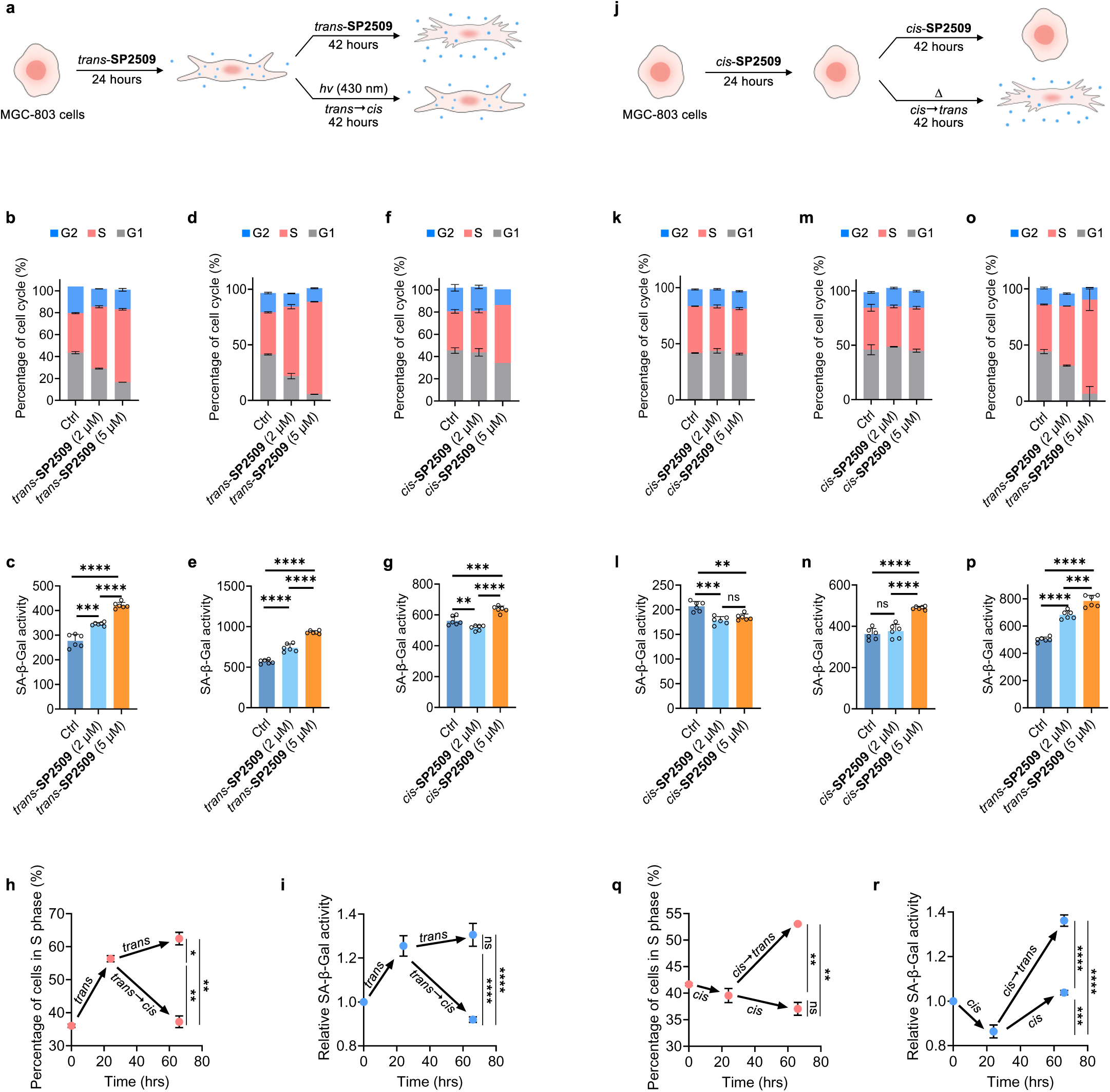
Optical switching of SP2509 dynamically controls senescence-associated trajectories in MGC-803 cells. **a**, Schematic of the *trans*-to-*cis* switching protocol. MGC-803 cells were treated with *trans*-**SP2509** in the dark for 24 hours and were then either maintained in the dark for a further 42 hours or transferred to 430-nm illumination to enrich the *cis*-state. **b**,**c**, Cell-cycle distribution (b) and senescence-associated β-galactosidase (SA-β-Gal) activity (c) after the initial 24-hour treatment with *trans*-**SP2509** at the indicated concentrations. **d**,**e**, Cell-cycle distribution (d) and SA-β-Gal activity (e) after continuous treatment with *trans*-**SP2509** in the dark for 66 hours. **f**,**g**, Cell-cycle distribution (f) and SA-β-Gal activity (g) after 24 hours of treatment with *trans*-**SP2509**, followed by 42 hours of 430-nm illumination to induce the *trans*→*cis* transition. **h**,**i**, Time-resolved changes in the percentage of cells in S phase (h) and relative SA-β-Gal activity (i) under continuous dark treatment or following *trans*→*cis* photoconversion at 24 hours. **j**, Schematic of the reciprocal *cis*-to-*trans* switching protocol. Cells were first treated for 24 hours with pre-irradiated, *cis*-enriched **SP2509** under 430-nm illumination and were then either maintained under illumination for a further 42 hours or transferred to the dark to allow thermal *cis*→*trans* relaxation. **k**,**l**, Cell-cycle distribution (k) and SA-β-Gal activity (l) after the initial 24-hour treatment under *cis*-enriching conditions. **m**,**n**, Cell-cycle distribution (m) and SA-β-Gal activity (n) after continuous maintenance under *cis*-enriching illumination for 66 hours. **o**,**p**, Cell-cycle distribution (o) and SA-β-Gal activity (p) after 24 hours under *cis*-enriching conditions followed by 42 hours in the dark to permit *cis*→*trans* relaxation. **q**,**r**, Time-resolved changes in the percentage of cells in S phase (q) and relative SA-β-Gal activity (r) under continuous cis-enriching illumination or following transfer to the dark at 24 hours. In the stacked cell-cycle plots, G1, S and G2 populations are shown in grey, salmon and blue, respectively. Ctrl denotes vehicle-treated cells. Data are presented as mean ±s.e.m. Statistical significance was determined using an unpaired two-tailed Student’s *t*-test. ns, not significant; \**P* < 0.05; \*\**P* < 0.01; \*\*\**P* < 0.001; \*\*\*\**P* < 0.0001.

The reciprocal experiment tested whether LSD1 inhibition could be imposed after an initial protected period (Fig. 3j and Supplementary Fig. S9, S11, S12). Cells were first cultured with 430 nm pre-irradiated **SP2509** under intermittent light for 24 hours, conditions that maintain the *cis*-enriched, attenuated state. These cells remained morphologically similar to controls and showed no substantial increase in senescence-associated β-galactosidase activity or S-phase accumulation (Fig. 3k, 3l). Cultures were then either kept under light or transferred to the dark, allowing thermal regeneration of the active *trans*-state (*cis→trans*). Light-maintained cultures remained close to controls (Fig. 3m, 3n), whereas dark transfer re-engaged senescence-associated β-galactosidase activity and S-phase accumulation (Fig. 3o-3r). Together with the reverse switch, this experiment shows that **SP2509** does not simply weaken senescence induction, but temporally gates entry into and exit from a plastic arrest trajectory.

### Photostate history controls population-level apoptotic commitment

We next examined whether the same timing principle extends to an effectively irreversible fate. In OCI-AML3 acute myeloid leukemia cells, Annexin V-FITC/propidium iodide (PI) flow cytometry showed that *trans*-**SP2509** increased apoptosis in a concentration-dependent manner after 48 hours (Fig. 4a,b and Supplementary Fig. S13). In the flow-cytometry plots, viable cells were mainly distributed in Q4, whereas apoptotic cells were quantified as the combined Annexin V-positive Q3 and Q2 populations, corresponding to early apoptosis and late apoptosis, respectively. Vehicle-treated cells were dominated by Q4 cells (85.3%) with only a small Q2/Q3 apoptotic fraction (14.65%), whereas *trans*-**SP2509** produced a concentration-dependent increase in the Q2 + Q3 apoptotic population across the tested dose range and at the highest dose, 10 µM of *trans*-**SP2509** shifted the population toward Q2/Q3 (84.5%) and reduced the Q4 fraction to 15.3%. On the other hand, at 0.5–2 µM, *cis*-enriched **SP2509** caused little apoptosis (Q2+Q3 14.2–15.6%) and largely preserved the Q4 population (85.8–87.1%), whereas at 5–10 µM it increased Q2/Q3 populations (26.9–51.3%) but less strongly than *trans*-**SP2509** at matched concentrations. The Q1 population remained comparatively minor across conditions (Supplementary Fig. S13), indicating that the dominant effect of **SP2509** photostate switching was regulation of apoptotic entry rather than primary necrotic death.

**Fig. 4.**
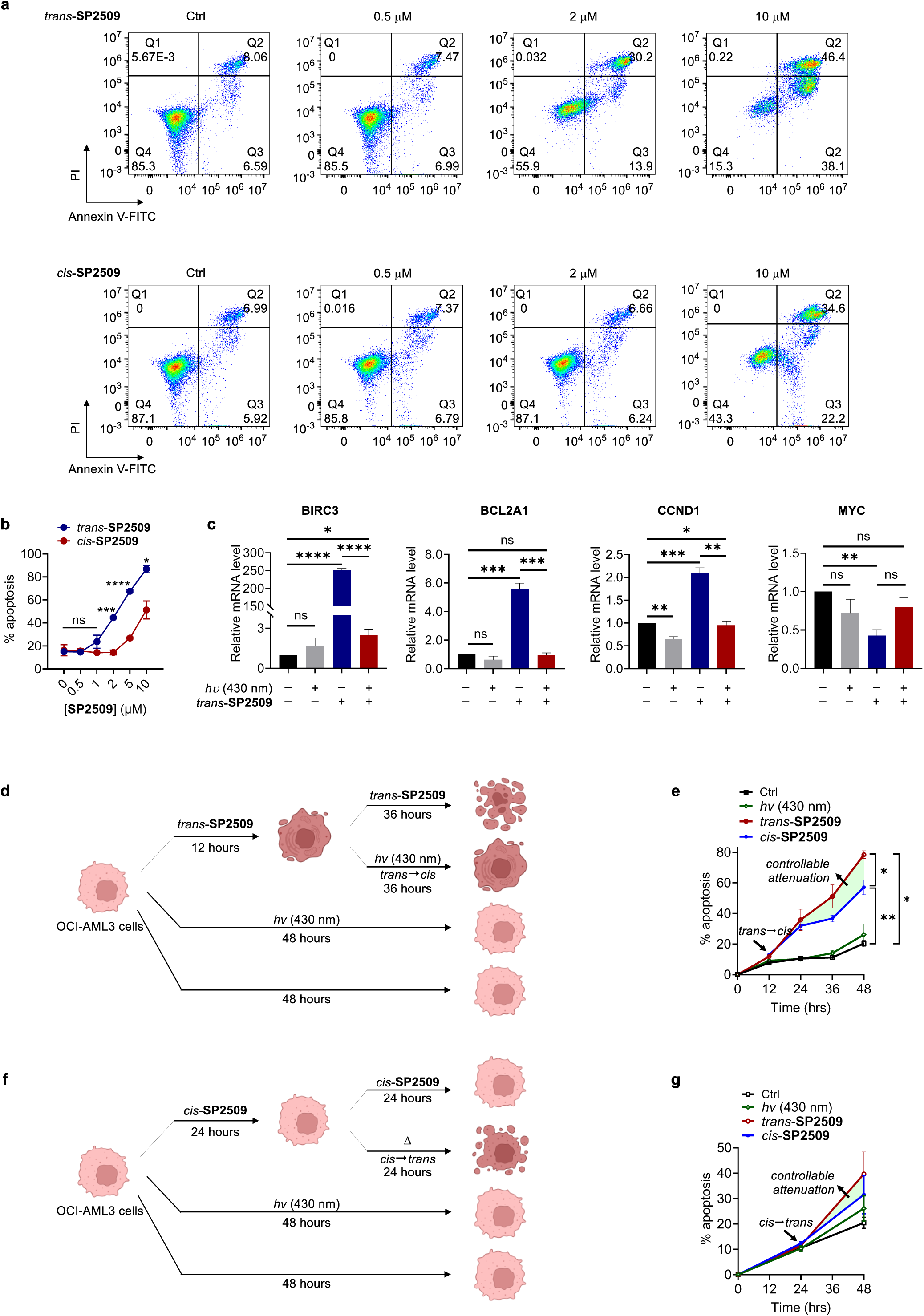
Optical switching of SP2509 controls population-level apoptotic commitment in OCI-AML3 cells. **a**, Representative Annexin V-FITC/propidium iodide flow-cytometry plots showing apoptosis in OCI-AML3 cells treated with increasing concentrations of *trans*-**SP2509** or *cis*-enriched **SP2509** for 48 hours. **b**, Quantification of apoptotic cells across **SP2509** concentrations, showing stronger apoptosis induction by *trans*-**SP2509** than by *cis*-enriched **SP2509**. **c**, Relative mRNA expression of apoptosis- and cell-cycle-associated genes after **SP2509** treatment. *trans*-**SP2509** strongly induced BIRC3, BCL2A1 and CCND1, whereas cells maintained under *cis*-enriched **SP2509** conditions showed attenuated responses; MYC displayed a distinct response pattern. **d**, Schematic of the *trans*-to-*cis* switching experiment. Cells were treated with 5 µM of *trans*-**SP2509** in the dark for 12 hours and then transferred to 430 nm illumination for 36 hours to enrich the *cis*-state. **e**, Time-resolved apoptosis analysis showing that photoconversion after 12 hours attenuated apoptotic accumulation compared with continuous dark treatment. **f**, Schematic of the reciprocal *cis*-to-*trans* switching experiment. Cells were first maintained under 430 nm illumination for 24 hours to keep **SP2509** in the *cis*-enriched attenuated state and were then transferred to the dark for 24 hours to allow thermal recovery of the active *trans*-state. **g**, Time-resolved apoptosis analysis showing increased apoptotic accumulation after transfer to the dark compared with continued illumination.

To probe the transcriptional basis of this differential survival response, we selected BIRC3, BCL2A1 and CCND1 as NF-κB-linked survival and cell-cycle regulators that are frequently engaged downstream of epigenetic stress adaptation. MYC was included as a central proliferation and metabolic regulator whose expression can be uncoupled from anti-apoptotic buffering programs (Fig. 4c). Quantitative reverse-transcription polymerase chain reaction (PCR) after 24 hours revealed strongly photostate-associated transcriptional responses. At 5 µM, *trans*-**SP2509** increased BIRC3 expression to 252-fold above vehicle, whereas *cis*-enriched **SP2509** induced only a low-amplitude response of 2-fold. The same photostate bias was observed for BCL2A1 and CCND1, with *trans*-**SP2509** increasing their expression to 5.6-fold and 2.1-fold, respectively, whereas the corresponding *cis*-enriched conditions remained close to baseline at 0.9-fold. By contrast, MYC showed a weaker and directionally distinct response, changing from 0.4-fold under *trans*-**SP2509** treatment to 0.8-fold under *cis*-enriched conditions (Fig. 4c). This pattern is consistent with a model in which *trans*-**SP2509** engages a compensatory stress-survival circuit that buffers proliferative drive while simultaneously priming apoptotic commitment. In contrast, cells maintained under *cis*-enriched **SP2509** conditions fail to robustly activate this adaptive transcriptional program.

To test dynamic control, OCI-AML3 cells were first treated with 5 µM of *trans*-**SP2509** in the dark for 12 hours and were then either maintained in the dark or transferred to 430 nm illumination to initiate the *trans*→*cis* transition for a further 36 hours (Fig. 4d and Supplementary Fig. S14, S15). Apoptosis levels were similar at the 12-hour transition point, with all treatment groups showing approximately 11.8–13.1% apoptotic cells (Fig. 4e), indicating comparable early engagement before trajectory divergence. Thereafter, continuous dark treatment drove a sustained increase in apoptosis, reaching approximately 35.8% at 24 hours, 51.1% at 36 hours and 78.4% at 48 hours. By contrast, switching toward the *cis*-enriched configuration produced a controllable attenuation of this trajectory, with the apoptotic fraction rising more slowly to approximately 31.9% at 24 hours, 36.7% at 36 hours and 57.1% at 48 hours (Fig. 4e and Supplementary Fig. S14, S15). Control and light-only groups remained substantially lower, reaching only approximately 20.4–26.1% apoptosis at 48 hours. Thus, even after initial exposure to the active inhibitor state, switching **SP2509** into the less active photostate does not abolish the early pro-apoptotic signal, but slows its subsequent population-level propagation toward apoptotic commitment.

In the reciprocal experiment, cells were first cultured with *cis*-enriched **SP2509** under 430 nm illumination for 24 hours and were then either maintained under light or transferred to the dark to allow thermal *cis*→*trans* relaxation (Fig. 4f). At the 24-hour transition point, apoptosis remained low and comparable across groups, at approximately 10.5–12.3% (Fig. 4g). By 48 hours, dark transfer increased the apoptotic fraction from approximately 31.6% under continuous illumination to 39.7%, corresponding to an approximately 26% relative increase upon *cis*→*trans* relaxation (Fig. 4g). This response is consistent with recovery of the more active *trans*-state after removal of 430 nm illumination. Control and light-only groups remained lower, at approximately 20.4–26.1 % apoptosis at 48 hours. This reciprocal switch shows that attenuation by the initial *cis*-enriched state is not fixed, but can be released when **SP2509** is allowed to thermally regenerate the active *trans*-state. Apoptosis remains irreversible after individual cells pass a commitment point, but these reciprocal switching experiments show that the population-level fraction entering apoptosis can be tuned through controllable attenuation and recovery of LSD1 inhibition.

### RNA-seq defines photostate-associated stress-survival circuitry

We used RNA-seq to define the transcriptional state that connects optical target control to apoptotic outcome. OCI-AML3 cells were treated for 24 hours under four matched conditions corresponding to the RNA-seq groups: control (Ctrl), light-only (*hν* (430 nm)), *trans*-**SP2509** and *cis*-**SP2509** (5 µM of **SP2509** under dark or 430 nm illumination, respectively) (Fig. 5). This design allowed us to separate transcriptional effects caused by light exposure alone from those associated with the active or attenuated **SP2509** photostate. Principal-component analysis separated *trans*-**SP2509** samples from control, light-only and *cis*-**SP2509** samples, with PC1 and PC2 explaining 28.4% and 18.5% of the variance, respectively (Fig. 5a). Sample-correlation analysis similarly showed that *trans*-**SP2509** formed a transcriptionally distinct group (Supplementary Fig. S16A). Thus, the more active *trans*-state does not merely increase the magnitude of a light-induced response, but establishes a separable transcriptional state.

**Fig. 5.**
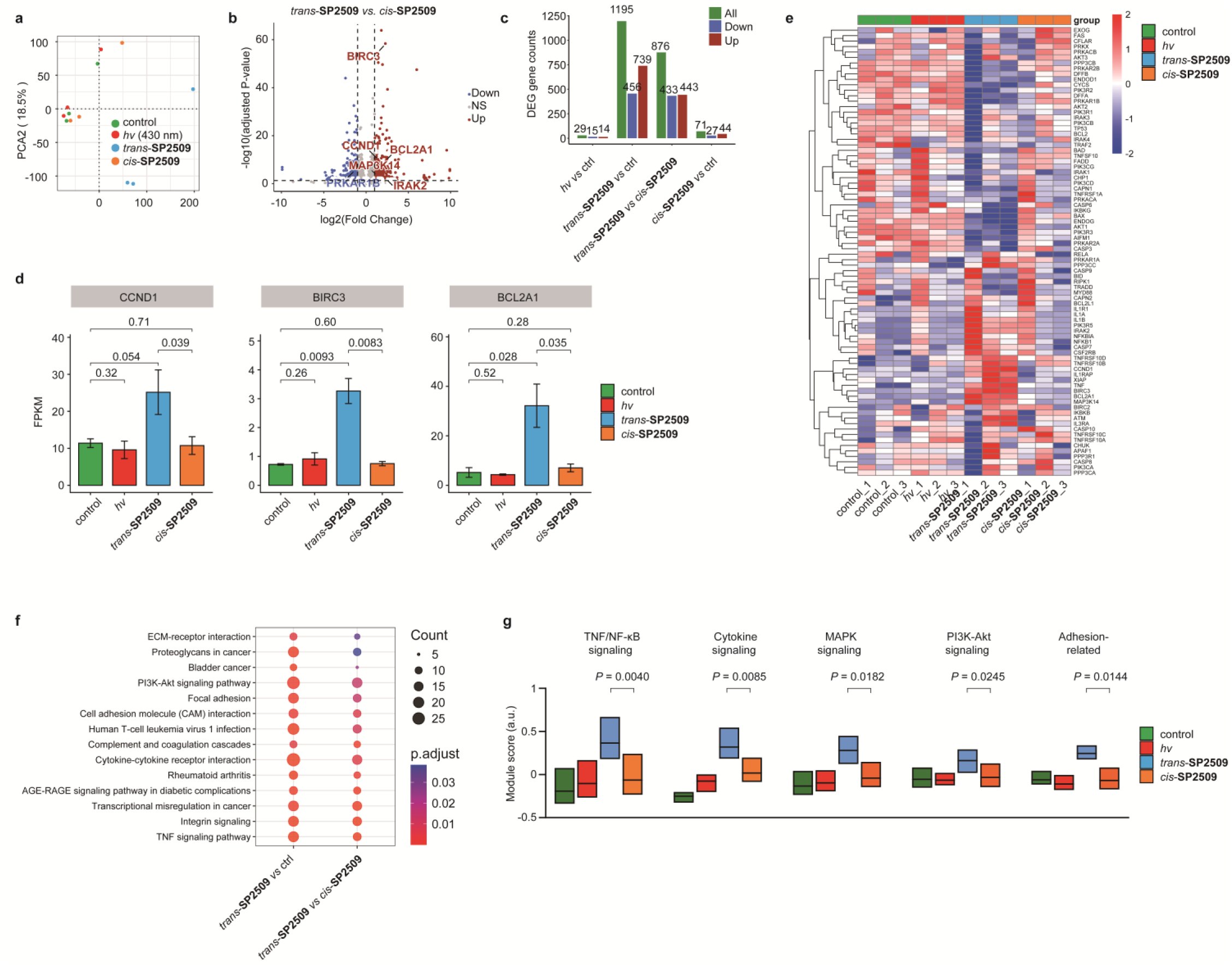
RNA-seq reveals photostate-dependent stress-survival transcriptional programs. a, Principal-component analysis of RNA-seq profiles from OCI-AML3 cells treated for 24 hours under four matched conditions: control, *hν* (430 nm), *trans*-**SP2509** and *cis*-**SP2509**. PC1 and PC2 explain 28.4% and 18.5% of the variance, respectively. **b,** Volcano plot comparing *trans*-**SP2509** and *cis*-**SP2509**. Genes with |log2(fold change)| ≥ 1 and adjusted P ≤ 0.05 are highlighted, and apoptosis-related genes are labelled. **c,** Differentially expressed gene counts across pairwise comparisons. *hν* alone induced few changes, whereas *trans*-**SP2509** produced a broad transcriptional response relative to control and to *cis*-**SP2509**. **d,** FPKM values of representative genes CCND1, BIRC3 and BCL2A1. *Trans*-**SP2509** induced higher expression of these stress-survival and cell-cycle-associated genes than control, *hν* or *cis*-**SP2509**. **e,** Heatmap of apoptosis-related genes across individual RNA-seq samples. *trans*-**SP2509** generated a distinct apoptosis- and stress-associated expression pattern that was attenuated in the *cis*-**SP2509** group. **f,** KEGG enrichment analysis of upregulated genes from the indicated comparisons. Enriched pathways include adhesion-related programs, PI3K-Akt signaling, cytokine-cytokine receptor interaction and TNF signaling. **g,** Module-score analysis of selected stress-survival pathways. *Trans*-**SP2509** produced higher pathway scores for TNF/NF-κB, cytokine, MAPK, PI3K-Akt and adhesion-related signaling than *cis*-**SP2509**.

Pairwise differential-expression analysis revealed that 430 nm illumination alone produced few changes, whereas *trans*-**SP2509** induced a broad gene-expression program relative to control (Fig. 5b,c and Supplementary Fig. S16B). Many of these changes were reduced in cells maintained under *cis*-**SP2509** conditions, indicating that the transcriptome responds to the photochemical state of the inhibitor rather than to illumination itself. The direct *trans*-versus *cis*-comparison highlighted apoptosis- and survival-associated genes, including BIRC3 and BCL2A1, which encode *anti*-apoptotic regulators commonly linked to NF-κB-dependent stress adaptation (Fig. 5c,d and Supplementary Fig. S16B). Consistent with quantitative reverse-transcription PCR (Fig. 4c), RNA-seq FPKM values for CCND1, BIRC3 and BCL2A1 were highest in cells treated with dark *trans*-**SP2509**, whereas cells maintained under *cis*-enriched **SP2509** conditions showed expression levels closer to control or light-only groups. These data suggest that active LSD1 inhibition does not simply trigger apoptosis, but also engages a compensatory survival program that accompanies apoptotic priming.

Heatmaps of apoptosis-related genes across individual samples and group averages showed that *trans*-**SP2509** selectively activated a stress- and survival-associated transcriptional module that was not reproduced by light alone or by the *cis*-enriched treatment condition (Fig. 5e-g and Supplementary Fig. S16C). KEGG analysis of upregulated genes identified enrichment in pathways associated with apoptosis, TNF/NF-κB signaling, cytokine-cytokine receptor interactions, MAPK signaling, PI3K-Akt signaling and adhesion-related programs (Fig. 5e,f and Supplementary Fig. S16D). These pathways are consistent with a mixed stress-adaptation state in which pro-death signaling, inflammatory-survival signaling and cell-state remodeling are activated together. Module-score analysis supported this photostate-associated pattern (Fig. 5g). Pathway-level scores were highest in the dark *trans*-**SP2509** group and lower in cells maintained under *cis*-**SP2509** conditions, indicating that photoconversion attenuates the coordinated pathway-level program rather than selectively suppressing a single downstream gene.

RNA-seq therefore links the photochemical state of **SP2509** to cell-fate outcome. In the more active *trans*-associated state, cells activate both pro-apoptotic signals and compensatory survival pathways, bringing the population closer to apoptotic commitment. By contrast, the *cis*-enriched state weakens this coordinated stress-response programme rather than simply producing a smaller version of the same response. This difference may explain why light can change the proportion of cells that enter apoptosis, even though apoptosis cannot be reversed once individual cells have committed to it.

### A chemical-to-cell-fate mechanism for optical epigenetic control

The combined data support a mechanism in which **SP2509** converts a photochemical change in hydrazone geometry into a timed perturbation of LSD1-dependent cell fate. At the molecular level, 430 nm illumination shifts **SP2509** from an extended *trans*-geometry that favors LSD1 inhibition toward a compact *cis*-enriched ensemble with weaker target engagement. At the chromatin level, this change reduces the effective strength of LSD1 inhibition and attenuates H3K9me2 accumulation. At the cellular level, the same photostate change is translated into altered senescence-associated arrest in MGC-803 cells and altered apoptotic entry in OCI-AML3 cells. This mechanism rationalizes the cell-type-specific outputs observed across the study. In MGC-803 cells, *trans*-**SP2509** initiates a senescence-associated trajectory marked by enlarged and flattened morphology, increased senescence-associated β-galactosidase activity and S-phase accumulation. Switching to the *cis*-enriched state after 24 hours attenuates this trajectory, revealing that the early arrest program remains reversible within the tested window. In OCI-AML3 cells, *trans*-**SP2509** activates a stress-adaptation transcriptional state that includes BIRC3, BCL2A1 and CCND1 together with TNF/NF-κB, cytokine, MAPK, PI3K-Akt and adhesion-linked pathway scores. Switching to, or maintaining cells in, the *cis*-enriched state reduces this stress-survival program and decreases the fraction of cells that proceed to apoptosis.

Thus, **SP2509** functions not as a lower fixed dose, but as a reversible timing device for LSD1 inhibition. A conventional dose reduction can reduce pathway amplitude, but it cannot alter inhibitor activity after a trajectory has begun. By contrast, photostate switching allows LSD1 inhibition to be attenuated or restored at defined times, thereby exposing reversible windows, commitment thresholds and delayed fate responses in living cells.

## DISCUSSION

This study identifies **SP2509** as a bioactive acylhydrazone whose intrinsic *trans*/*cis* photoisomerization can be harnessed to control LSD1-dependent biology. The advance is not the installation of an external photoswitch into an inhibitor, but the discovery of a latent switch within a pharmacophore already optimized for biological activity. DFT optimization of the deprotonated phenolate forms showed that *cis*-**SP2509** adopts a compact, intramolecularly hydrogen-bonded pseudocycle, providing a plausible structural rationale for its attenuated LSD1 inhibition relative to the open *trans*-conformation. The same change in hydrazone geometry is transmitted to recombinant enzyme inhibition, chromatin methylation, proliferation, senescence, apoptosis and transcriptional state.

Several features make **SP2509** well suited to optical control of chromatin. First, the more active *trans*-state is regenerated thermally in the dark, allowing inhibitory activity to recover without adding a second reagent. Second, 430 nm light creates a less active *cis*-state under physiological conditions and under low-intensity intermittent illumination in cells. Third, the biochemical switching is large enough to create clear phenotypic differences but not so absolute that dose-dependent partial inhibition is lost. This behavior is useful for probing threshold effects in chromatin signaling, where intermediate enzyme activity can yield outcomes that differ qualitatively from complete inhibition.

The work also provides a biological lesson. LSD1 inhibition is often treated as a static perturbation of chromatin, but the switching experiments show that the order and duration of inhibition matter. Senescence-associated phenotypes in MGC-803 cells were reversible or re-engageable within the tested window, whereas apoptosis in OCI-AML3 cells behaved as an irreversible single-cell fate whose population frequency could nevertheless be tuned by light. These observations separate target reversibility from cellular irreversibility and show how a photoswitchable epigenetic inhibitor can map the temporal logic of cell-fate decisions.

The RNA-seq data support a chromatin-linked transcriptional mechanism in which hydrazone isomerization changes LSD1 engagement and thereby modulates stress-survival circuitry. Dark *trans*-**SP2509** induces apoptosis-associated and compensatory survival genes, including BIRC3, BCL2A1 and CCND1, together with pathway-level enrichment in TNF/NF-κB, cytokine, MAPK, PI3K-Akt and adhesion-related signaling. Cells maintained under *cis*-**SP2509** conditions show weaker engagement of these programs. This result suggests that LSD1 inhibition does not simply trigger a single linear death pathway; rather, it creates a stress-adaptation state in which survival and death programs compete, and the photostate of **SP2509** controls how rapidly the population moves through this state. More broadly, these findings suggest that latent photoswitches in existing chemical probes and drug-like molecules can provide a route to photo-pharmacology without rebuilding the lead scaffold. Acylhydrazones are well represented among bioactive small molecules, but their isomerization behavior is often not analyzed during biological evaluation. **SP2509** shows that such hidden dynamics can be converted from a potential complication into a useful control principle. The same logic may apply to other scaffolds in which a dynamic covalent or proton-coupled motif is already positioned within a target-binding pharmacophore.

Further work should measure photo-stationary distributions and thermal relaxation directly in live-cell media, quantify cellular LSD1 engagement with orthogonal assays and map genome-wide chromatin changes at LSD1-bound loci. These studies will clarify how much of the observed transcriptional response is driven by local LSD1 occupancy, by secondary chromatin adaptation or by stress-response amplification. Even with these open questions, **SP2509** provides both a tool for temporally controlled epigenetic perturbation and a design principle for discovering optically tunable activity in established bioactive chemotypes.

## METHODS

### General information

Unless otherwise stated, reagents were obtained from commercial suppliers and used without further purification. ^1^H and ^13^C NMR spectra were recorded on Bruker 300, 400 or 500 MHz spectrometers. Chemical shifts are reported in ppm relative to residual solvent signals. High-resolution mass spectra were obtained by electrospray ionization mass spectrometry.

### Synthesis of *trans*-SP2509

*trans*-**SP2509** was synthesized from 3-(chlorosulfonyl)benzoic acid through morpholine sulfonamide formation, methyl esterification, hydrazide formation and condensation with 1-(5-chloro-2-hydroxyphenyl)ethanone. The final product was isolated by filtration and crystallization from petroleum ether/ethyl acetate (4/1) and was obtained as a solid in 32% yield over the final two steps. Analytical data were consistent with the reported structure, and full procedures and spectra are provided in the Supplementary Information.^12^

### Computational analysis

Geometry optimizations and harmonic frequency calculations were performed using Gaussian 09, Revision D.01, at the ωB97X-D/6-31G(d) level. Solvation in DMSO was described using the integral-equation-formalism polarizable continuum model (IEFPCM). The deprotonated phenolate forms of *trans*- and *cis*-**SP2509** were treated as closed-shell singlet anions with a total charge of −1 and a multiplicity of 1. The initial *cis*-geometry was generated by rotating the N–N=C–Ar dihedral of the *trans*-structure towards 0°, followed by unconstrained geometry optimization. Harmonic frequency calculations were performed at the same level of theory and confirmed that both optimized structures were local minima with no imaginary frequencies. Thermochemical corrections were evaluated at 298.15 K and 1 atm. Selected geometrical parameters were calculated from the final optimized Cartesian coordinates. Complete Gaussian input and output files, optimized Cartesian coordinates, frequency-analysis results and thermochemical quantities are provided in the Supplementary Information and Supplementary Data.

### NMR analysis of SP2509 photoisomerization

*trans*-**SP2509** (5 mg) was dissolved in DMSO-*d*_6_ (0.6 ml) and transferred to an NMR tube. After acquisition of the initial spectrum, triethylamine (4 equiv.) was added directly to the NMR tube. The sample was irradiated with a 430 nm lamp (20 mW·cm^−2^) for 20 minutes, followed by 365 nm irradiation (70 mW·cm^−2^) for 20 minutes. The sample was then incubated in the dark for 12 hours. ^1^H NMR spectra were acquired after each step.

### UV-vis spectroscopy

**SP2509** was prepared as a 50 µM solution in DMSO/H_2_O (1:9, *V*/*V*) or DMSO/PBS (1/9, *V*/*V*; 1×PBS, pH 7.4). UV-vis spectra were recorded before and after 430 nm irradiation. For photoconversion experiments, the PBS solution was irradiated at 430 nm (2 mW·cm^−2^) for 10, 40 or 60 minutes. For dark-relaxation experiments, the *cis*-enriched solution was kept at room temperature in the dark and spectra were recorded over time. Recovery of absorbance at 415 nm was fitted to estimate the thermal half-life.

### Recombinant LSD1 inhibition assay

LSD1 inhibition was measured using a Cayman Chemical LSD1 screening kit (#700120), which couples LSD1-dependent H_2_O_2_ formation to horseradish peroxidase-mediated oxidation of ADHP to resorufin. *trans*-**SP2509** samples containing 3 equiv. DBU were diluted to 20×final concentrations in DMSO and divided into dark and irradiated aliquots. Irradiated aliquots were exposed to 430 nm light (5 mW·cm^−2^) for 45 minutes before addition to the 96-well assay plate. Fluorescence was measured according to the manufacturer’s protocol and dose-response curves were fitted to determine IC_50_ values.

### Cell culture and illumination protocols

MGC-803 and HeLa cells were cultured in high-glucose DMEM supplemented with 10% fetal bovine serum and 1% penicillin/streptomycin. OCI-AML3 cells were cultured in IMEM supplemented with 10% fetal bovine serum and 1% penicillin/streptomycin. Cells were maintained at 37 °C under 5% CO_2_. For light-treated groups, media containing **SP2509** were pre-irradiated at 430 nm, typically 5 mW·cm^−2^ for 30 minutes, to enrich the *cis* state and then applied to cells. During incubation, cells were intermittently irradiated with low-intensity 430 nm light using the duty cycles indicated in the figure legends and Supplementary Information.

### H3K9me2 Simple Western assay

HeLa cells were treated with **SP2509** under dark or 430 nm illuminated conditions for 48 hours. Cells were lysed in nuclear extraction buffer, histones were acid-extracted and the supernatant was analyzed on a ProteinSimple Jess system. H3K9me2 was detected with an anti-H3K9me2 antibody and normalized to total H3. Signal linearity and antibody dilutions were optimized before comparison of treatment conditions.

### Cell viability and senescence assays

For cell viability analysis, MGC-803 cells were treated with serial concentrations of *trans*-**SP2509** or *cis*-enriched **SP2509** for 48 hours and viability was determined using a CCK-8 assay. Senescence-associated β-galactosidase staining was performed using a Solarbio SA-β-gal staining kit (#G1580). Cells were fixed and stained at 37 °C overnight, imaged by microscopy and quantified using a fluorescence-based SA-β-gal activity assay as described in the Supplementary Information.

### Flow cytometry

Apoptosis in OCI-AML3 cells was measured using Annexin V-FITC and propidium iodide staining followed by analysis on a FACSCalibur flow cytometer. FITC-positive events were counted as apoptotic cells, and the apoptotic fraction was calculated as the sum of the Q2 and Q3 populations. Cell-cycle distributions in MGC-803 cells were measured after ethanol fixation, RNase A treatment and propidium iodide staining using a Solarbio DNA content quantitation assay (#CA1510). Flow cytometry data were analyzed using FlowJo 10.8.

### Quantitative reverse-transcription PCR

OCI-AML3 cells were treated with 5 µM of **SP2509** under dark or 430 nm illuminated conditions for 24 hours. Total RNA was isolated, reverse-transcribed and analysed using TaqMan probes for BIRC3, BCL2A1, CCND1 and MYC. GAPDH was used for normalization. Primer and probe sequences are listed in Supplementary Table 1.

### RNA sequencing and analysis

OCI-AML3 cells were treated for 24 hours with DMSO or 5 µM **SP2509** under dark or 430 nm illuminated conditions, with three biological replicates per group. Cell pellets were frozen in liquid nitrogen and sent to Beijing Novogene Co., Ltd. for sequencing. Libraries were prepared with the Fast RNA-seq Lib Prep Kit V2 (ABclonal, RK20306) and sequenced on an Illumina NovaSeq X Plus platform to generate 150 bp paired-end reads. Reads were aligned to the human GRCh38 primary assembly using HISAT2. Differential expression was analysed using DESeq2. Genes with absolute log2(fold change) >1 and adjusted P ≤ 0.05 were considered differentially expressed, and GO and KEGG enrichment analyses were performed using clusterProfiler.^40–42^

### Statistics

Data are presented as mean ±s.e.m. or mean ±s.d. as indicated in the figure legends. Statistical significance was assessed using unpaired two-tailed Student’s *t*-tests unless otherwise indicated. Exact replicate numbers and *P* values should be added to the final source-data tables before submission.

## Data availability

RNA-seq data have been deposited in the Human Research Archive under accession code HRA019264, associated with BioProject accession PRJCA067391. Source data for graphs and uncropped or raw Simple Western and flow-cytometry files will be provided with the final submission.

## ACKNOWLEDGEMENTS

This work was supported by the National Key R&D Program of China (2025YFA0920900), Strategic Priority Research Program of the Chinese Academy of Sciences (XDB0960103), Beijing National Laboratory for Molecular Sciences (BNLMS-CXTD-202202), the Open Fund of the State Key Laboratory of Elemento-Organic Chemistry, Nankai University (202206), and the National Natural Science Foundation of China (22537005, 22271291, and 22527901 through the National Major Research Instrumentation Program). We would like to thank Prof. Peng-Xu Qian at Zhejiang University for kindly providing the OCI-AML3 cells, Dr. Kun Pang and Prof. Jun-Chao Shi at China National Center for Bioinformation/Beijing Institute of Genomics, Chinese Academy of Sciences for performing RNA-seq analysis, and Prof. Xiao-Bo Qiu at Beijing Normal University for insightful discussions. C.-S. W. acknowledges the support of a BMS Junior Fellowship from the Beijing National Laboratory for Molecular Sciences (BNLMS).

## Author contributions

CW: Data curation, Formal Analysis, Investigation, Software, Supervision, Validation, Visualization; LL: Funding acquisition; LC: Conceptualization, Data curation, Formal Analysis, Funding acquisition, Project administration, Resources, Supervision, Visualization, Writing – original draft, Writing – review & editing.

