## Supplemental Information for "A latent hydrazone switch turns SP2509 into an optical timer of cell fate"

**Table of contents**

Procedures for the Synthesis of *trans*-**SP2509** S2

Computational analysis of deprotonated **SP2509** S3

NMR Spectral Characterization of *trans*-**SP2509** Isomerization S7

UV-Vis Characterization of *trans*-**SP2509** Isomerization S7

LSD1 Inhibition Assay S8

Cell Viability Assay S9

Detection of H3K9me2/H3 by Simple Western blot S9

Senescence-Associated β-Galactosidase Staining S10

Quantification of Senescence-Associated β-Galactosidase activity S11

Flow Cytometric Detection of Cell Cycle S14

Quantitative Reverse Transcription PCR S17

Flow Cytometric Detection of Apoptosis S18

RNA-seq Analysis S22

References S24

NMR Spectrum S25

**General Information**

Unless otherwise noted, all reagents were obtained from commercial suppliers and used without further purification. $^{1}H$ NMR and $^{13}C$ NMR spectra were recorded on Bruker 300, 400 or 500 MHz spectrometers. Chemical shifts (δ) are reported in ppm relative to the residual solvent signals for $^{1}H$ and $^{13}C$ NMR ($CDCl_3$: $^{1}H$ NMR, 7.26 ppm; $^{13}C$ NMR, 77.0 ppm). Multiplicities are reported as follows: singlet (s), doublet (d), doublet of doublets (dd), triplet (t), quartet (q) and multiplet (m). HRMS data were obtained by electrospray ionization mass spectrometry (ESI-MS; Thermo Fisher Scientific).

**Procedures for the Synthesis of *trans*-SP2509** [1]

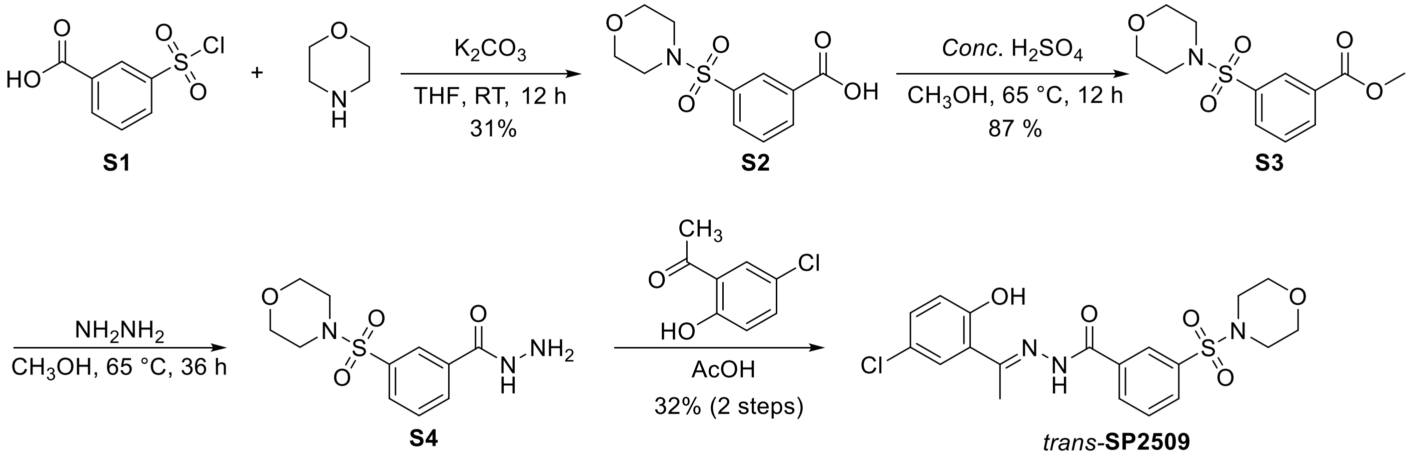


**Fig. S1.** Synthesis of *trans*-**SP2509**.

To a solution of morpholine (13.1 mmol, 1.18 mL) in tetrahydrofuran (40 mL) were added potassium carbonate (22.67 mmol, 3.14 g) and 3-(chlorosulfonyl)benzoic acid **S1** (9.07 mmol, 2.0 g). The resulting reaction mixture was stirred at room temperature for 12 hours. After completion of the reaction, the solvent was removed under reduced pressure. The crude material was washed with 1 M HCl (25 mL × 2) and extracted with dichloromethane. The organic layer was dried over sodium sulfate and concentrated under reduced pressure. The residue was purified by silica gel column chromatography to afford **S2** (750 mg, 31%) as a white solid. $^{1}H$ NMR (400 MHz, DMSO-$d_6$): δ 13.61 (br, 1H), 8.28 (d, $J$ = 7.8 Hz, 2H), 8.20 (s, 2H), 7.99 (d, $J$ = 7.9 Hz, 2H), 7.83 (d, $J$ = 7.8 Hz, 1H), 3.69-3.59 (m, 6H), 2.96-2.83 (m, 6H). $^{13}C$ NMR (100 MHz, DMSO-$d_6$): δ 166.0, 135.0, 133.9, 132.2, 131.6, 130.2, 128.0, 65.3, 45.8. HRMS calculated for $C_{11}H_{12}NO_5S$ $[M-H]^-$: 270.0442, found: 270.0438.

To a solution of **S2** (2.77 mmol, 750 mg) in methanol (30 mL) was slowly added concentrated sulfuric acid (2.77 mmol, 1.5 mL). The resulting reaction mixture was heated at 65 °C for 12 hours. After completion of the reaction, the solvent was removed under reduced pressure, and the crude material was purified by silica gel column chromatography to afford **S3** (689 mg, 87%) as a white solid. $^{1}H$ NMR (400 MHz, $CDCl_3$): δ 8.39 (s, 1H), 8.29 (d, $J$ = 7.8 Hz, 1H), 7.94 (d, $J$ = 7.8 Hz, 1H), 7.66 (t, $J$ = 7.8 Hz, 1H), 3.96 (s, 3H), 3.82-3.65 (m,

4H), 3.16-2.91 (m, 4H). $^{13}$C NMR (100 MHz, $CDCl_3$): δ 165.4, 136.0, 133.9, 131.8, 131.4, 129.4, 128.8, 66.0, 52.7, 46.0. HRMS calculated for $C_{12}H_{16}NO_5S$ $[M+H]^+$: 286.0744, found: 286.0739.

To a solution of **S3** (2.42 mmol, 689 mg) in methanol (30 mL) was slowly added hydrazine hydrate (13.31 mmol, 0.9 mL). The resulting reaction mixture was heated at 65 °C for 36 hours. After completion of the reaction, the solvent was removed under reduced pressure, and the crude material was used in the next step without further purification. $^1$H NMR (400 MHz, DMSO-$d_6$): δ 10.10 (s, 1H), 8.18-8.13 (m, 2H), 7.87 (d, $J$ = 7.9 Hz, 1H), 7.76 (t, $J$ = 7.7 Hz, 1H), 4.66 (s, 2H), 3.65-3.61 (m, 4H), 2.91-2.84 (m, 4H). $^{13}$C NMR (100 MHz, DMSO-$d_6$): δ 164.2, 134.8, 134.4, 131.7, 130.0, 129.8, 126.0, 65.2, 45.9. HRMS calculated for $C_{11}H_{14}N_3O_4S$ $[M-H]^-$: 284.0711, found: 284.0706.

Under a nitrogen atmosphere, 1-(5-chloro-2-hydroxyphenyl)ethanone (1 mmol, 170 mg) and acetic acid (0.5 mmol, 31.5 µL) were added to a solution of **S4** (1 mmol, 285 mg) in methanol (35 mL). The resulting reaction mixture was heated at 120 °C for 72 hours. After completion of the reaction, *trans*-**SP2509** was obtained by filtration and crystallization from a mixed solvent system (petroleum ether/ethyl acetate = 4/1) to afford a solid product (140 mg, 32%). $^1$H NMR (400 MHz, DMSO-$d_6$): δ 13.33 (s, 1H), 11.70 (s, 1H), 8.29 (d, $J$ = 7.8 Hz, 1H), 8.20 (s, 1H), 7.98 (d, $J$ = 7.9 Hz, 1H), 7.85 (t, $J$ = 7.8 Hz, 1H), 7.67 (d, $J$ = 2.1 Hz, 1H), 7.36 (dd, $J$ = 8.8, 2.1 Hz, 1H), 6.96 (d, $J$ = 8.8 Hz, 1H), 3.69-3.59 (m, 4H), 2.97-2.87 (m, 4H), 2.50 (s, 3H). HRMS calculated for $C_{19}H_{19}N_5O_3ClS$ $[M-H]^-$: 436.0739, found: 436.0734.

**Computational analysis of deprotonated SP2509**

Geometry optimizations and harmonic frequency calculations were performed using Gaussian 09, Revision D.01. The deprotonated phenolate forms of *trans*- and *cis*-**SP2509** were treated as closed-shell singlet anions with a total charge of −1 and a multiplicity of 1. All calculations were carried out using the long-range-corrected ωB97X-D density functional and the 6-31G(d) basis set. Solvation in DMSO was described using the integral-equation-formalism polarizable continuum model (IEFPCM). The Gaussian route section was:

```
# opt freq wb97xd/6-31g(d) scrf=(iefpcm,solvent=dmso)
```

The initial *trans*-structure was constructed with an extended hydrazone geometry in which the N–N=C–Ar dihedral was close to 180°. The initial *cis*-structure was generated from the optimized *trans* geometry by rotating the N–N=C–Ar dihedral towards 0° and orienting the phenolate oxygen towards the hydrazide N–H group. Both structures were subsequently optimized without geometrical constraints at the same level of theory.

Geometry optimization converged for both isomers. Harmonic frequency calculations performed at the corresponding optimized geometries gave no imaginary frequencies (NImag = 0), confirming that both structures correspond to local minima on the potential-energy surface. The lowest calculated vibrational frequencies were 8.7038 $cm^{-1}$ for *trans*-**SP2509** and 14.4522 $cm^{-1}$ for *cis*-**SP2509**. The final electronic energies were −2133.82954503 Hartree for *trans*-**SP2509** and −2133.85324338 Hartree for *cis*-**SP2509**. Both the optimization and frequency calculations terminated normally.

Selected geometrical parameters were obtained directly from the final optimized Cartesian coordinates. Atom numbering followed that used in the Gaussian input and output files. The hydrazone dihedral was defined as D(N9–N14–C15–C17), where N9 is the hydrazide nitrogen bearing H34, N14 is the imine nitrogen, C15 is the imine carbon and C17 is the ipso carbon of the aryl group attached to the C=N bond. The intramolecular hydrogen-bond parameters were defined using the phenolate oxygen O24, the hydrazide hydrogen H34 and its donor nitrogen N9. Accordingly, the reported hydrogen-bond distances correspond to O24···H34 and O24···N9, and the donor–hydrogen–acceptor angle corresponds to ∠N9–H34···O24. The sign of a dihedral angle depends on the specified atom order and coordinate orientation; absolute dihedral values were therefore used when comparing the extent of *trans* and *cis* geometry in the main text and figures.

The optimized *trans*-structure retained an open conformation, with D(N9–N14–C15–C17) = −178.088°. The phenolate oxygen remained remote from the hydrazide N–H group, giving O24···H34 and O24···N9 distances of 4.287 and 4.129 Å, respectively, and an ∠N9–H34···O24 angle of 74.25°. In contrast, *cis*-**SP2509** retained a near-zero hydrazone dihedral of 2.413° and adopted a folded pseudocyclic conformation. This geometry supported an intramolecular phenolate O···H–N hydrogen bond, with O24···H34 and O24···N9 distances of 1.577 and 2.581 Å, respectively, and an ∠N9–H34···O24 angle of 158.02°.

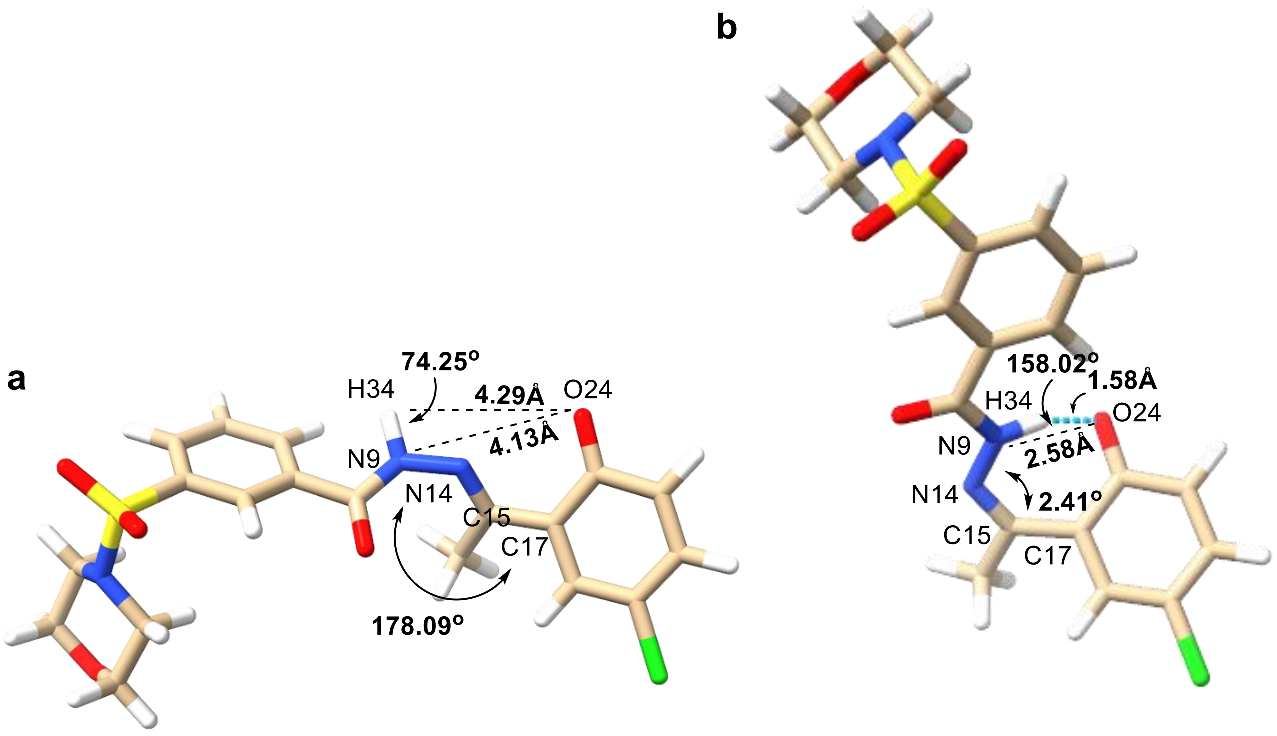


**Supplementary Fig. S2. DFT-optimized geometries and selected structural parameters of deprotonated *trans*- and *cis*-SP2509**. **a,** Front view of optimized *trans*-**SP2509**, showing an open conformation with an absolute N9–N14–C15–C17 dihedral angle of 178.09°. The phenolate oxygen remained remote from the hydrazide N–H group, with O24···H34 and O24···N9 distances of 4.29 and 4.13 Å, respectively, and an N9–H34···O24 angle of 74.25°. **b,** Corresponding front view of optimized *cis*-**SP2509**. The *cis* isomer retained an absolute N9–N14–C15–C17 dihedral angle of 2.41° and adopted a pseudocyclic conformation stabilized by an intramolecular phenolate O24···H34–N9 hydrogen bond. The O24···H34 and O24···N9 distances were 1.58 and 2.58 Å, respectively, and the N9–H34···O24 angle was 158.02°. Structures were optimized at the ωB97X-D/6-31G(d) level using IEFPCM solvation in DMSO. The figures of the structural models were prepared in ChimeraX[2].

**Supplementary Table S1. Selected calculated properties of deprotonated *trans*- and *cis*-SP2509**

| **Parameter** | ***trans*-SP2509** | ***cis*-SP2509** |
|---|---|---|
| Total charge | −1 | −1 |
| Multiplicity | 1 | 1 |
| Electronic energy, Hartree | −2133.82954503 | −2133.85324338 |
| D(N9–N14–C15–C17), ° | −178.088 | 2.413 |
| \|D(N9–N14–C15–C17)\|, ° | 178.088 | 2.413 |

| Parameter | *trans*-SP2509 | *cis*-SP2509 |
|---|---|---|
| O24···H34 distance, Å | 4.287 | 1.577 |
| O24···N9 distance, Å | 4.129 | 2.581 |
| N9–H34 distance, Å | 1.012 | 1.050 |
| ∠N9–H34···O24, ° | 74.25 | 158.02 |
| Lowest vibrational frequency, $cm^{-1}$ | 8.7038 | 14.4522 |
| Number of imaginary frequencies | 0 | 0 |

Thermochemical corrections were obtained from the harmonic frequency calculations using the Gaussian default temperature and pressure of 298.15 K and 1 atm. The resulting electronic, zero-point-corrected, thermal, enthalpic and Gibbs free-energy quantities are reported for computational transparency. These quantities describe individual optimized conformers of a single deprotonated microstate within a harmonic-frequency and continuum-solvent approximation. The calculations do not include explicit solvent molecules, exhaustive conformational populations, anharmonic or hindered-rotor corrections, alternative protonation or tautomeric states, electronically excited states, or the kinetic barriers connecting the two isomers. Consequently, the calculated energy difference between the two optimized structures should not be interpreted as an experimental *trans*/*cis*-isomerization free energy, a prediction of the photostationary distribution, or a direct measure of the thermal relaxation rate. The calculations were used primarily to compare the intrinsic geometrical organization of the two deprotonated photostates.

**Supplementary Table S2. Calculated electronic energies and thermochemical corrections for deprotonated *trans*- and *cis*-SP2509.** Geometry optimizations and harmonic frequency calculations were performed at the ωB97X-D/6-31G(d) level using IEFPCM solvation in DMSO. All values are reported in Hartree.

| Quantity | *trans*-SP2509 | *cis*-SP2509 |
|---|---|---|
| SCF electronic energy | −2133.82954503 | −2133.85324338 |
| Zero-point energy correction | 0.372280 | 0.371495 |
| Thermal correction to energy | 0.398066 | 0.397205 |
| Thermal correction to enthalpy | 0.399011 | 0.398149 |
| Thermal correction to Gibbs free energy | 0.312967 | 0.312833 |
| Electronic energy + zero-point energy | −2133.457265 | −2133.481748 |
| Electronic energy + thermal Gibbs free energy | −2133.516578 | −2133.540410 |

**NMR Spectral Characterization of *trans*-SP2509 Isomerization**

The tautomerization process of *trans*-**SP2509** was characterized by $^1$H NMR spectroscopy. *trans*-**SP2509** (5 mg) was added to a 1.5 mL centrifuge tube and dissolved in DMSO-$d_6$ (0.6 mL). The solution was transferred to an NMR tube, and the initial $^1$H NMR spectrum was acquired. After acquisition, triethylamine (4 equiv.) was added directly to the NMR tube, and a second $^1$H NMR spectrum was recorded. The NMR tube was then irradiated with a 430 nm lamp (20 mW·cm$^{-2}$) for 20 minutes, after which another $^1$H NMR spectrum was acquired. The same sample was subsequently irradiated with a 365 nm lamp (70 mW·cm$^{-2}$) for 20 minutes, followed by acquisition of another $^1$H NMR spectrum. Finally, the NMR tube was kept in the dark for 12 hours, and the final $^1$H NMR spectrum was recorded.

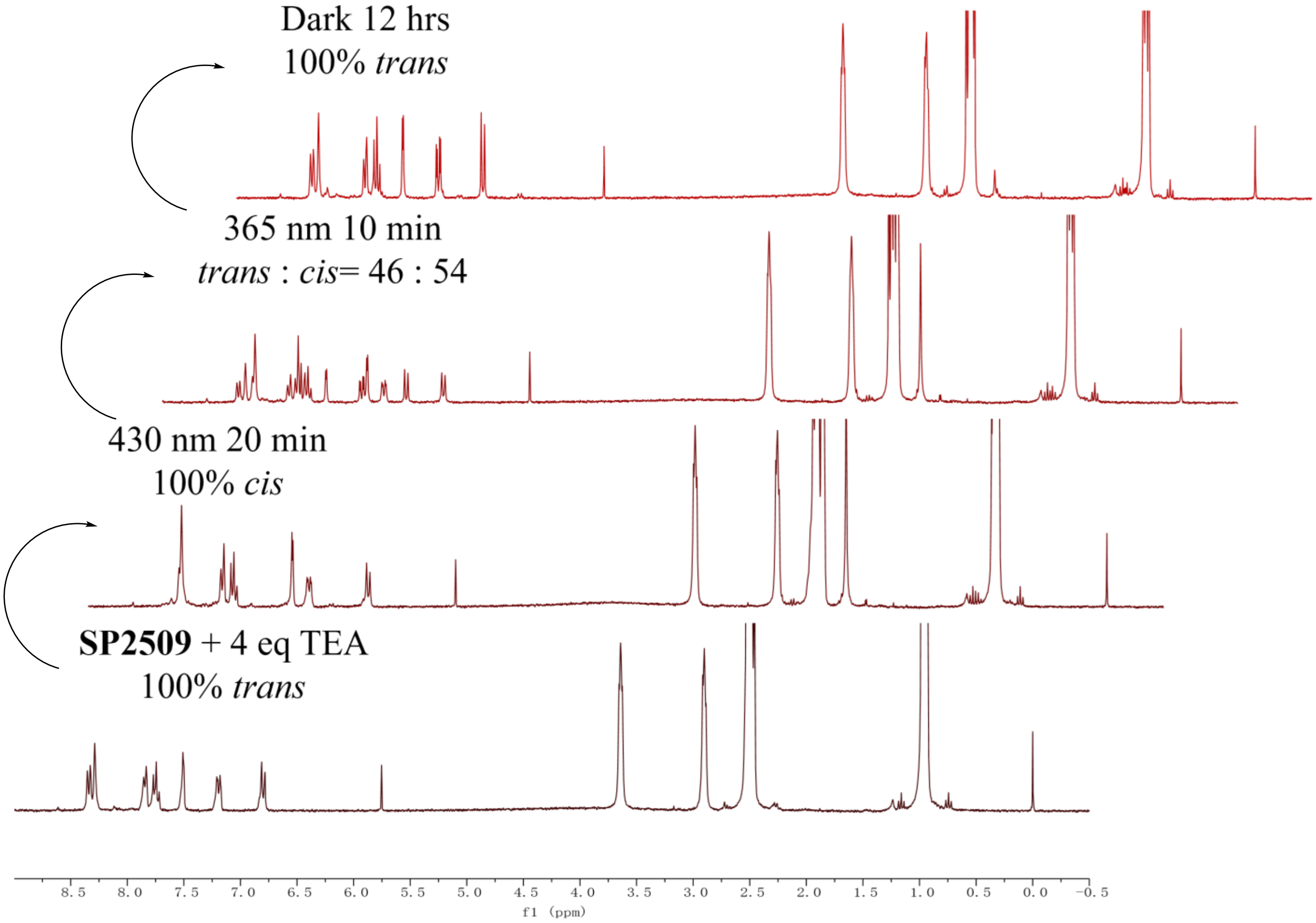


**Fig. S3.** NMR characterization of **SP2509** isomerization.

**UV-Vis Characterization of *trans*-SP2509 Isomerization**

A 50 µM solution of *trans*-**SP2509** was prepared in DMSO/$H_2O$ (1:9, v/v), and a second 50 µM solution was prepared in DMSO/PBS (1:9, v/v; Gibco, 1×, pH 7.4). UV-Vis absorption spectra of both solutions were recorded from 250 to 800 nm using a Lambda 950 UV-Vis spectrophotometer (PerkinElmer). To characterize *trans*-**SP2509** photoisomerization, the 50 µM *trans*-**SP2509** solution in DMSO/PBS (1:9, v/v; Gibco, 1×, pH

7.4) was irradiated at 430 nm (2 mW·cm$^{-2}$) for 10, 40 or 60 minutes. The solution reached the photostationary state after 40 minutes of irradiation. Additional irradiation for 20 minutes produced no further change in the UV-Vis absorption spectrum. After the photostationary state had been reached, the solution was stored in the dark at room temperature, and UV-Vis spectra were recorded at 5, 10, 20, 30, 45, 60, 90 minutes and at 2, 3, 4, 5, 6, 7 and 8 hours. The half-life of *cis*-**SP2509** was obtained by fitting the time-dependent absorbance at 415 nm using GraphPad Prism 8.4.

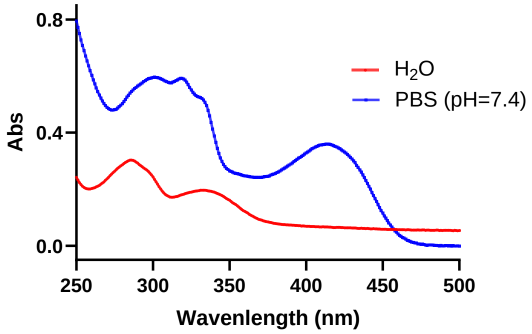


**Fig. S4.** The UV-Vis absorption spectrum of *trans*-**SP2509** in $H_2O$ and PBS solution

**LSD1 Inhibition Assay**

The LSD1 inhibitor screening assay kit was purchased from Cayman Chemical (#700120). The assay is based on a multistep enzymatic reaction in which LSD1 produces $H_2O_2$ during demethylation of lysine 4 on a peptide corresponding to the first 21 amino acids of the histone H3 *N*-terminal tail. In the presence of horseradish peroxidase (HRP), $H_2O_2$ reacts with ADHP (10-acetyl-3,7-dihydroxyphenoxazine) to produce the highly fluorescent product resorufin. Resorufin fluorescence was measured using an excitation wavelength of 530-540 nm and an emission wavelength of 585-595 nm.

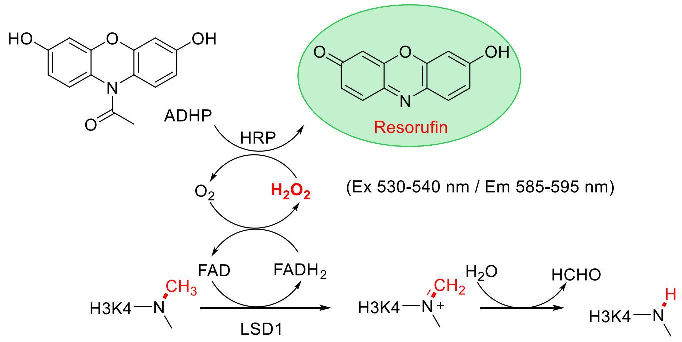


**Fig. S5.** The scheme of LSD1 inhibitor screening assay

*Trans*-**SP2509** samples containing 3 equiv. of 1,8-diazabicyclo(5.4.0)undec-7-ene (DBU) were diluted to 20× the desired final concentrations in 100% DMSO and divided into two aliquots. One aliquot (2.5 µL) was added directly to the 96-well plate, whereas the second aliquot was irradiated at 430 nm (5 mW·cm$^{-2}$) for 45 minutes (to convert *trans*-**SP2509** to *cis*-**SP2509**) before addition to the plate. The LSD1 enzyme stock was diluted 17-fold with assay buffer, and diluted LSD1 enzyme (40 µL) was added to the appropriate wells. The substrate mixture, consisting of horseradish peroxidase (3 μL), dimethyl-K4 peptide corresponding to the first 21 amino acids of the histone H3 *N*-terminal tail (3 μL), and ADHP (1.5 μL), was then added to each well. Resorufin fluorescence was measured on a microplate reader (BioTek Synergy H1) with excitation at 530 nm and emission at 590 nm. $IC_{50}$ values were obtained using GraphPad Prism 8.4.

**Cell Viability Assay**

MGC-803 cells were maintained in high-glucose Dulbecco's modified Eagle medium (DMEM; Gibco) supplemented with 10% fetal bovine serum (FBS; Gibco) and 1% penicillin/streptomycin (Gibco) at 37 °C under 5% $CO_2$. The CCK-8 cell proliferation and cytotoxicity assay kit (CA1210) was purchased from Beijing Solarbio Science & Technology Co., Ltd. MGC-803 cells were seeded into two 96-well plates at a density of $5 \times 10^3$ cells per well in 100 μL DMEM according to the manufacturer's instructions. After 24 hours, one plate was treated with fresh medium containing *trans*-**SP2509** at the indicated concentrations (0, 30 nM, 100 nM, 300 nM, 1 μM, 3 μM, 10 μM and 30 μM; 100 μL; 1% DMSO) and incubated in the dark for 48 hours at 37 °C. For the light-treated plate, media containing the corresponding concentrations of *trans*-**SP2509** were irradiated at 430 nm (5 $mW \cdot cm^{-2}$) for 30 minutes (to convert *trans*-**SP2509** to *cis*-**SP2509**) and then transferred to the cells. The light-treated plate was exposed intermittently to 430 nm light for 5 minutes every 30 minutes and incubated for 48 hours at 37 °C. After treatment, cells were washed twice with 1× PBS, and 100 μL fresh medium containing 10% CCK-8 reagent was added to each well. The plates were incubated for 30 minutes at 37 °C, and absorbance at 450 nm was measured using a microplate reader (BioTek Synergy H1).

**Detection of H3K9me2/H3 by Simple Western blot**

HeLa cells were maintained in high-glucose DMEM (Gibco) supplemented with 10% FBS (Gibco) and 1% penicillin/streptomycin (Gibco) at 37 °C under 5% $CO_2$. HeLa cells were seeded into six 6 cm cell culture dishes at a density of $1 \times 10^6$ cells per dish. After 12 hours, *trans*-**SP2509** at the indicated concentrations (0, 3 or 10 μM) was added to three dishes, which were then incubated in the dark for 48 hours at 37 °C. For the light-treated groups, media containing *trans*-**SP2509** (0, 3 or 10 μM) were pre-irradiated at 430 nm (5 $mW \cdot cm^{-2}$) for 30 minutes (to convert it to *cis*-**SP2509**) and then transferred to the remaining three dishes. These dishes were intermittently irradiated with 430 nm light (0.5 $mW \cdot cm^{-2}$; 10 seconds every 200 seconds) and incubated for 48 hours at 37 °C.

After the indicated treatments, histones were extracted from HeLa cells. Cells were harvested at the indicated time points and lysed on ice for 30 min in 1 mL nuclear extraction buffer (10 mM Tris-HCl, 10 mM $MgCl_2$, 25 mM KCl, 1% Triton X-100, 8.6% sucrose and protease inhibitor cocktail (Sangon, C600386)). Nuclei were collected by centrifugation at 1,000g for 5 minutes at 4 °C. The supernatant was removed, and histones were extracted for 4 hours with 0.4 N cold sulfuric acid (400 μL). Extracts were clarified by centrifugation at 15,000g for 5 minutes at 4 °C and transferred to fresh microcentrifuge tubes. 100% TCA solution (133 μL) was added to each tube. Histones were precipitated at 4 °C overnight, pelleted by centrifugation at 15,000g for

5 minutes, washed twice with cold acetone and resuspended in water. Histones were quantified using the BCA protein assay (Sangon, C503051). The H3K9me2/H3 ratio was then measured by Simple Western analysis.

The Simple Western assay was performed according to the manufacturer's protocol (ProteinSimple, Jess). To determine suitable sample concentrations and antibody dilutions, a preliminary experiment was performed using samples with known protein concentrations. Standard curves were established between the H3K9me2 NIR chemiluminescence signal area and total protein concentration, as well as between the H3 IR chemiluminescence signal area and total protein concentration. A linear relationship was observed for H3K9me2 and H3 signals at an antibody dilution of 1:20 and a total protein concentration of 0.1-0.4 mg·mL$^{-1}$. Therefore, 0.2 mg·mL$^{-1}$ was selected as the sample detection concentration, and 1:20 was selected as the antibody dilution.

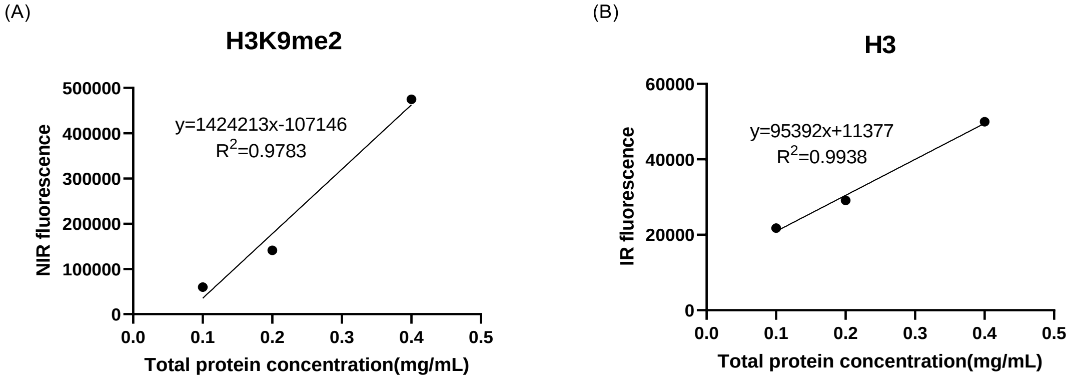


**Fig. S6.** Standard curves of IR and NIR signals as a function of total protein concentration.

After suitable sample concentrations and antibody dilutions had been determined, H3K9me2 levels were compared across the different treatment conditions using H3 as the loading control. Extracted histone solutions were diluted with 1× sample buffer and 5× master mix to a final protein concentration of 0.2 mg·mL$^{-1}$. The 12-230 kDa fluorescence separation module (SM-FL001) was used. H3K9me2 was detected with mouse anti-histone H3 antibody (Abcam, ab1220), and H3 was detected with rabbit anti-histone H3 antibody (Abcam, ab1791), each at an antibody dilution of 1:20. Anti-mouse NIR and anti-rabbit IR detection modules were used according to the manufacturer's instructions. Compass software (ProteinSimple) was used to visualize virtual gels. Relative protein quantities were generated from chromatograms of the indicated samples and corrected using the standard curves.

**Senescence-Associated β-Galactosidase Staining**

Senescence-associated β-galactosidase staining was performed using a SA-β-gal staining kit (Solarbio, #G1580). MGC-803 cells were seeded into four 6 cm cell culture dishes at a density of 5 × 10$^5$ cells per dish according to the manufacturer's instructions. After 24 hours, *trans*-**SP2509** at the indicated concentration (0 or 5 μM) was added to two dishes, which were then incubated in the dark for 48 hours at 37 °C. For the light-treated groups, media containing *trans*-**SP2509** (0 or 5 μM) were pre-irradiated at 430 nm (5 $mW \cdot cm^{-2}$) for 30 minutes (to convert *trans*-**SP2509** to *cis*-**SP2509**) and then transferred to the remaining two dishes. These dishes were intermittently irradiated with 430 nm light (0.5 $mW \cdot cm^{-2}$; 20 seconds every 100 seconds) and incubated for 48 hours at 37 °C. After incubation, cells were rinsed with phosphate-buffered saline, fixed and stained with SA-β-gal solution at 37 °C overnight. Cells were imaged using a microscope.

**Quantification of Senescence-Associated β-Galactosidase activity**

The senescence-associated β-galactosidase (SA-β-gal) assay was performed based on the protocol described in reference [3] with minor modifications. MGC-803 cells were maintained in high-glucose DMEM (Gibco) supplemented with 10% FBS (Gibco) and 1% penicillin/streptomycin (Gibco) at 37 °C under 5% $CO_2$. Briefly, MGC-803 cells were seeded at a density of $5 \times 10^5$ cells per well into two separate 6-well plates. After 12 h of incubation, the cells were treated with 0, 5, or 10 μM of *trans*-**SP2509**. The two plates were then divided into two experimental groups: one plate was incubated in the dark, and the other was exposed to light, both for 48 hours. For the light-treated groups, media containing *trans*-**SP2509** (0, 5 or 10 μM) were pre-irradiated at 430 nm (5 $mW \cdot cm^{-2}$) for 30 minutes (to convert *trans*-**SP2509** to *cis*-**SP2509**) and then transferred to the remaining plate. After incubation, cells were washed twice with PBS, harvested by trypsin digestion, and then lysed on ice for 10 min by the addition of 1 mL of 1X lysis buffer (5 mM 3-[(3-cholamidopropyl) dimethylammonio]-1-propanesulfonate [CHAPS], 40 mM citric acid, 40 mM sodium phosphate, 1X protease inhibitor cocktail, pH 6.0). The lysate was then centrifuged at 15,000 rpm for 10 minutes at 4 °C, and the supernatant was collected for further analysis. Reaction buffer at 2X strength consisted of 40 mM citric acid, 40 mM sodium phosphate, 300 mM NaCl, 10 mM β-mercaptoethanol, and 4 mM $MgCl_2$ (pH 6.0) with 1.7 mM of 4-methylumbelliferyl-β-D-galactopyranoside (MUG) added immediately prior to use from a 34 mM stock in dimethyl sulfoxide. An aliquot of 50 μL of cell lysate was mixed with 50 μL of 2X assay buffer in a 96-well plate. The mixture was incubated at 37 °C for 1 hour. After the reaction, 50 μL of the mixture was transferred to a black 96-well plate, and the reaction was terminated by adding 200 μL of 400 mM sodium carbonate aqueous solution. The fluorescence intensity was measured using a microplate reader with excitation at 360 nm and emission at 465 nm. The protein concentration of the cell lysate was determined by the BCA method. The normalized SA-β-gal activity was calculated as the fluorescence intensity divided by the protein concentration (ug/mL) and further divided by the reaction time (1 hour).

The experimental procedure for optically controlled dynamic regulation of senescence by **SP2509** is as follows: MGC-803 cells were seeded into 6 cm cell culture dishes at a density of $1 \times 10^5$ cells per dish. After 12 hours, *trans*-**SP2509** at the indicated concentrations (0, 2 or 5 μM) was added to the dishes.

For the *trans*-to-*cis* switching experiment, the dishes were divided into three groups. Group 1 was incubated in the dark for 24 hours, and SA-β-gal activity was then quantified as described above. Group 2 was first kept in the dark for 24 hours, then transferred to light for 42 hours; the light-transfer step consisted of continuous irradiation at 430 nm (5 mW·cm$^{-2}$) for 30 minutes, followed by intermittent irradiation at 430 nm (0.5 mW·cm$^{-2}$; 10 seconds every 200 seconds). After the total 66-hour incubation, SA-β-gal activity was quantified. Group 3 was maintained in the dark for 66 hours, and SA-β-gal activity was then determined.

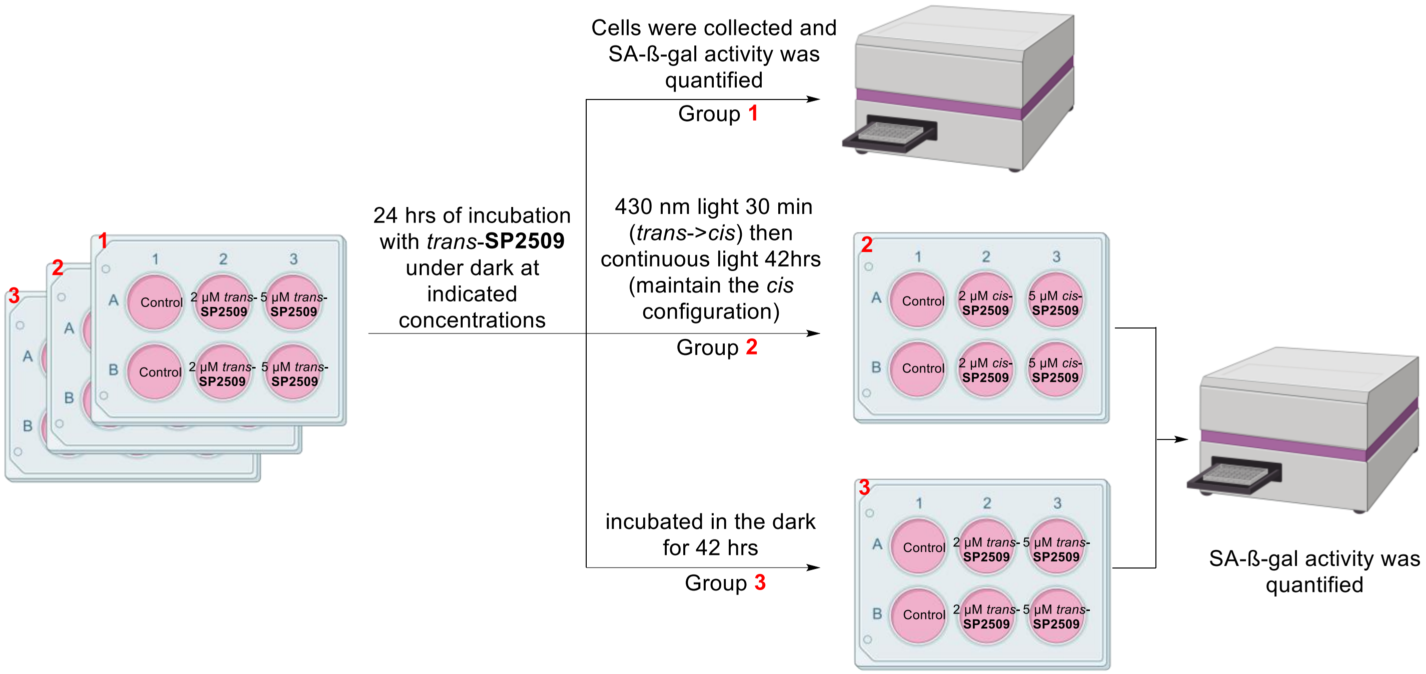


**Fig. S7.** Schematic of the *trans*-to-*cis* switching protocol and SA-β-gal detection.

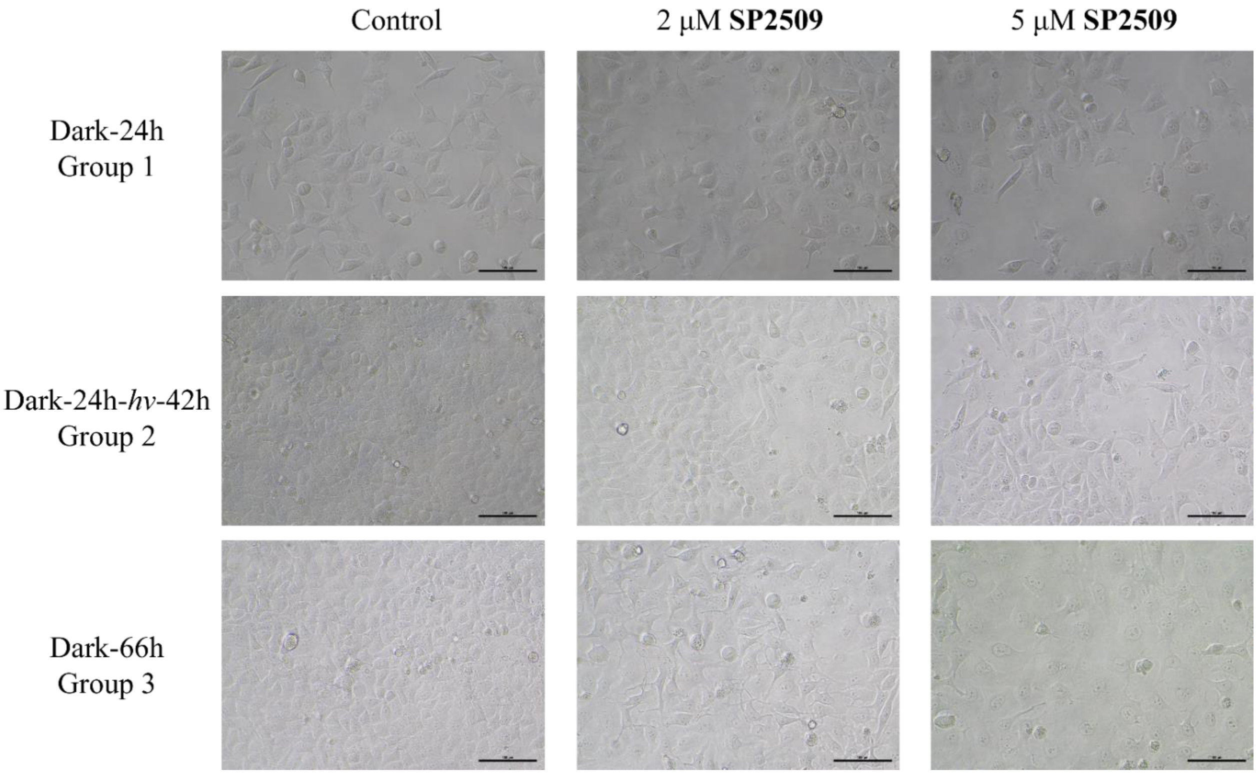


**Fig. S8.** Microscopic images of cells in response to *trans*-**SP2509** and photo-induced *trans*-to-*cis* isomerization.

For the *cis*-to-*trans* switching experiment, media containing *trans*-**SP2509** (0, 2, or 5 μM) were pre-irradiated at 430 nm (5 mW·cm$^{-2}$) for 30 minutes to convert the compound to the *cis*-isomer, and then transferred to the cells in fresh 6-cm dishes. The dishes were again divided into three groups. Group 1 was exposed to intermittent irradiation at 430 nm (0.5 mW·cm$^{-2}$; 10 seconds every 200 seconds) for 24 hours, after which SA-β-gal activity was measured. Group 2 was irradiated under the same intermittent condition for 24 hours and then transferred to the dark for an additional 42 hours, followed by quantification of SA-β-gal activity. Group 3 was maintained under continuous intermittent light for 66 hours, and SA-β-gal activity was then assessed.

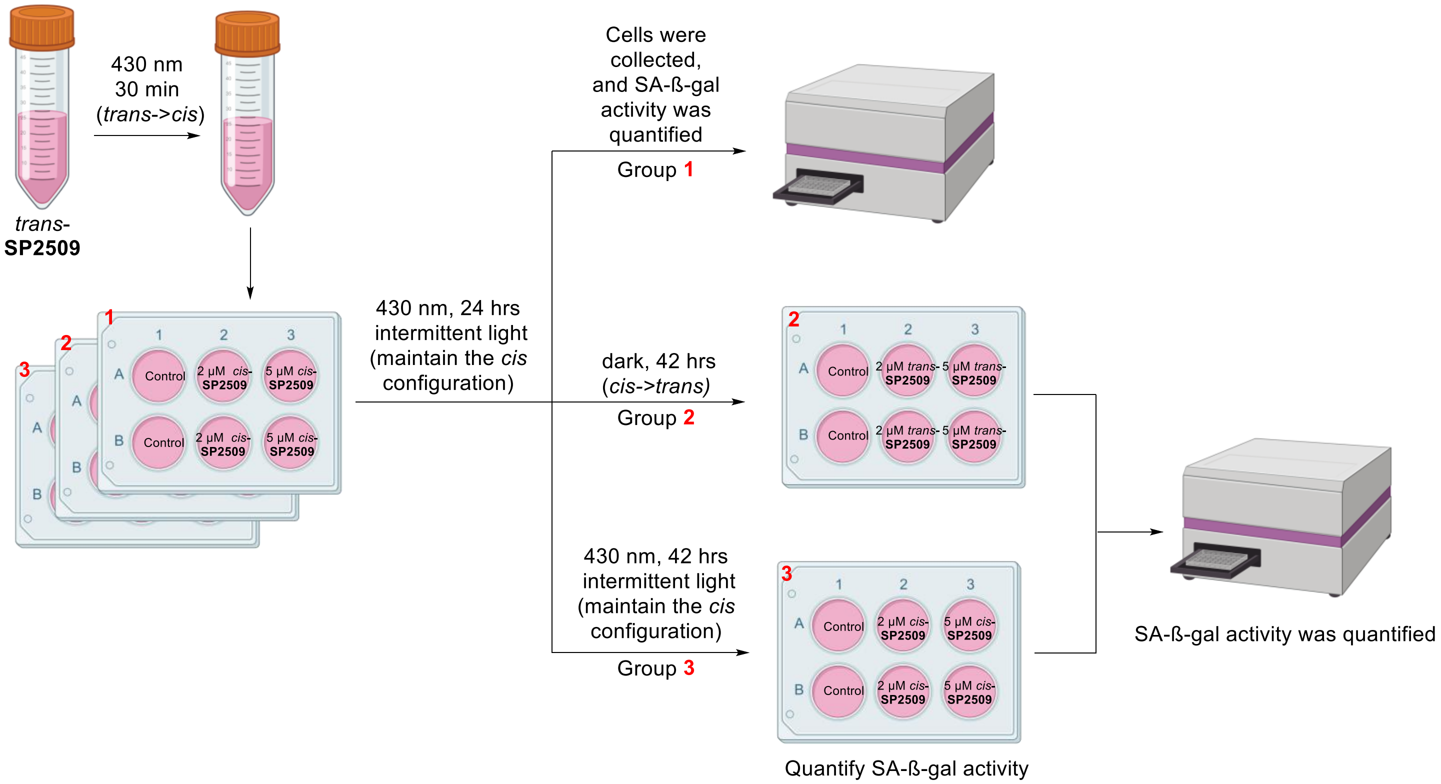


**Fig. S9.** Schematic of the reciprocal *cis*-to-*trans* switching protocol and SA-β-gal detection.

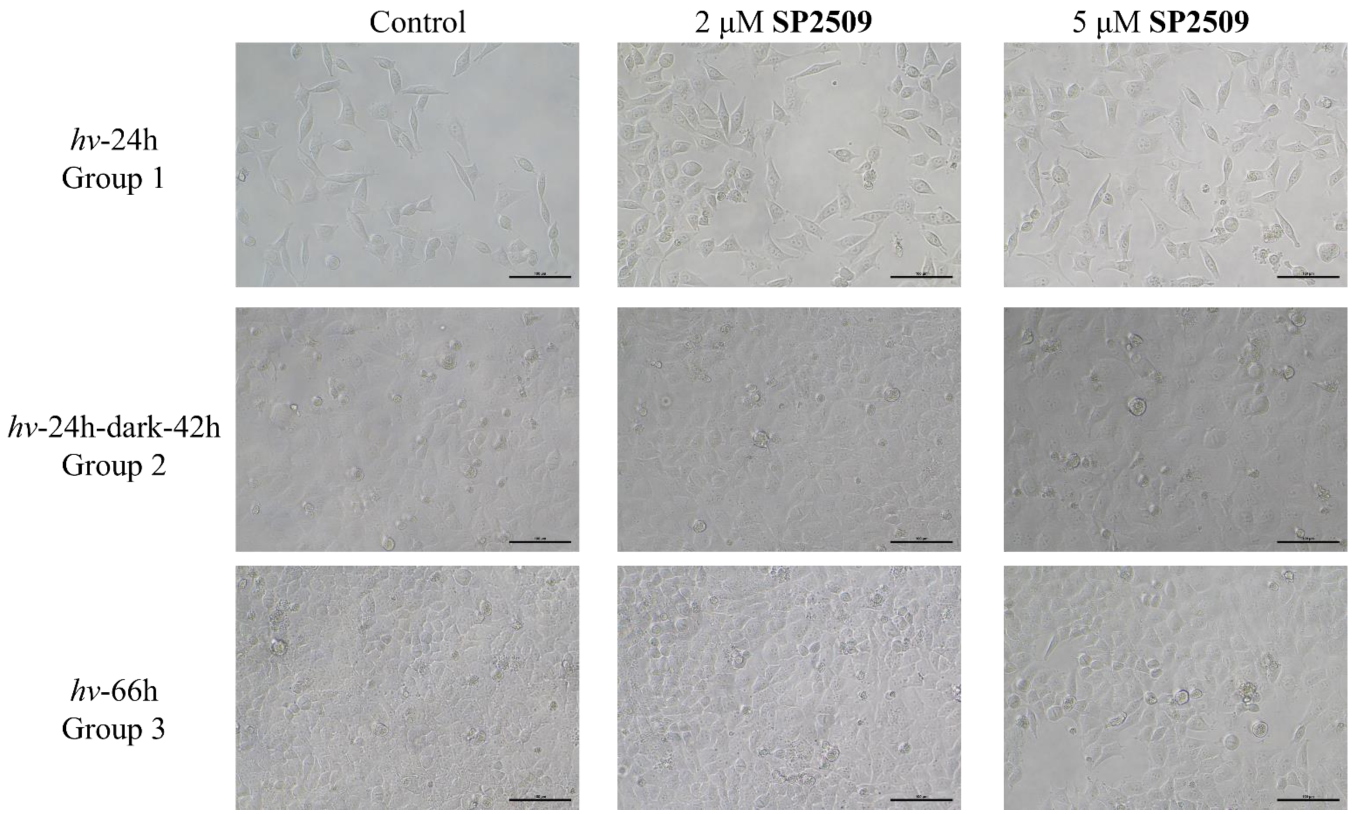


**Fig. S10.** Microscopic images of cells in response to *cis*-**SP2509**, and *cis*-to-*trans* isomerization.

**Flow Cytometric Detection of Cell Cycle**

MGC-803 cells were maintained in high-glucose DMEM (Gibco) supplemented with 10% FBS (Gibco) and 1% penicillin/streptomycin (Gibco) at 37 °C under 5% $CO_2$. MGC-803 cells were seeded into 6 cm cell culture dishes at a density of $5 \times 10^5$ cells per dish. After 12 hours, *trans*-**SP2509** at the indicated concentrations (0, 2 or 5 μM) was added to the dishes. For the *trans*-to-*cis* switching experiment, dishes were divided into three groups. The first group was incubated in the dark for 24 hours and then analyzed for cell cycle by flow cytometry. The second group was incubated in the dark for 24 hours and then transferred to light for 42 hours. The light-transfer step consisted of continuous irradiation at 430 nm (5 $mW \cdot cm^{-2}$) for 30 min followed by intermittent irradiation at 430 nm (0.5 $mW \cdot cm^{-2}$; 10 seconds every 200 seconds). The third group was maintained in the dark for 66 hours. Cell-cycle analysis was performed using a DNA content quantitation assay (Solarbio, #CA1510). Cells were collected, washed once with PBS and fixed in pre-cooled 70% ethanol for 6 hours at 4 °C. Fixed cells were centrifuged at 1,500 rpm for 5 minutes, washed once with PBS and incubated with 100 μL RNase A for 30 minutes at 37 °C. PI staining solution (400 μL) was added, and cells were incubated for 30 minutes at 4 °C in the dark. PI fluorescence was detected by flow cytometry, and the results were analyzed using FlowJo 10.8 software.

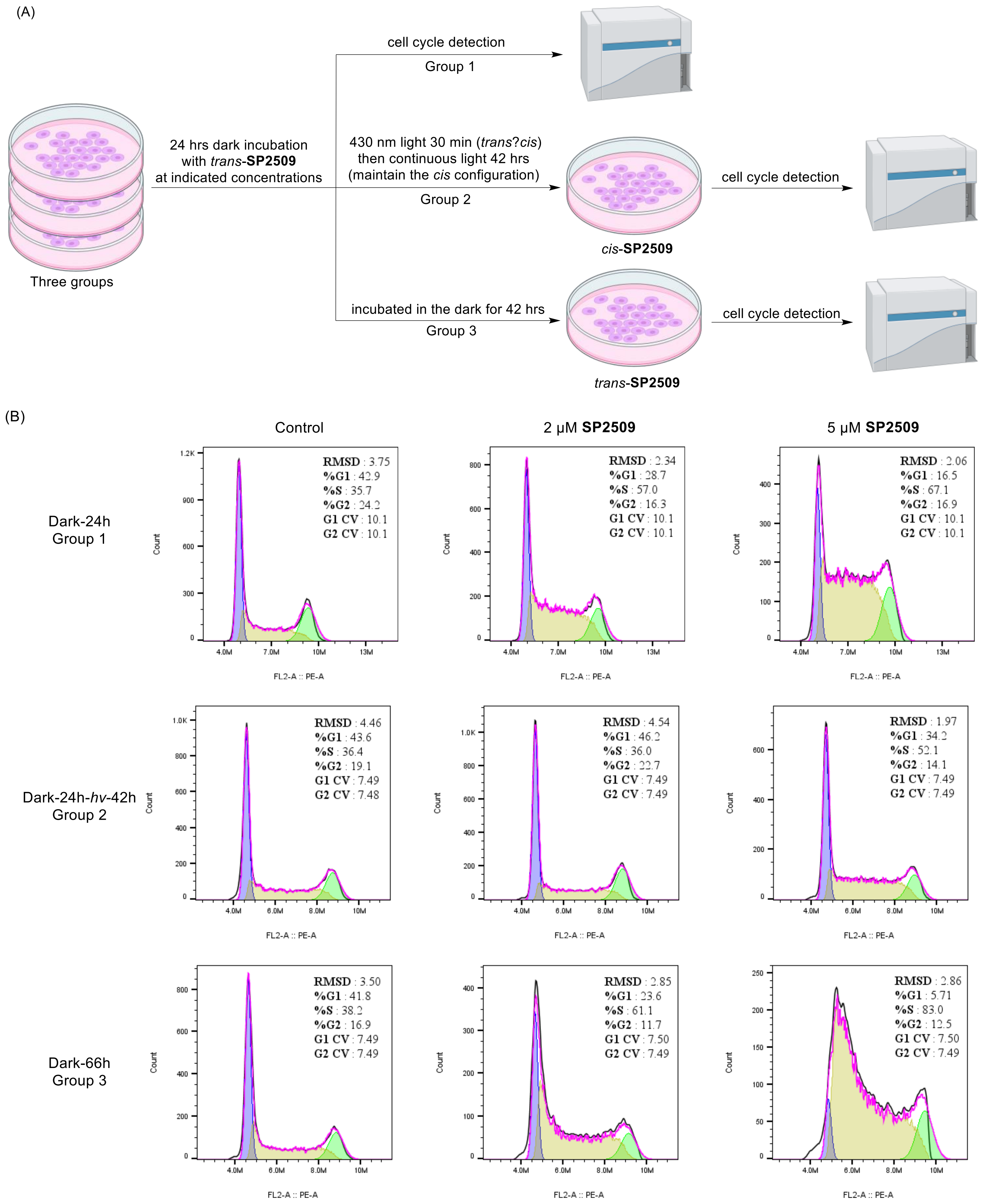


**Fig. S11.** (A) Schematic of the *trans*-to-*cis* switching protocol and cell cycle detection. (B) Cell cycle in response to *trans*-**SP2509**, and photo-induced *trans*-to-*cis* isomerization. The proportion of MGC-803 cells arrested in S phase increases after 24 hours of *trans*-**SP2509** treatment, returns to control levels upon light-driven *trans*-to-*cis* conversion for an additional 42 hours, and remains elevated after 66 hours of continuous *trans*-**SP2509** exposure.

For the *cis*-to-*trans* switching experiment, media containing *trans*-**SP2509** (0, 2 or 5 μM) were pre-irradiated at 430 nm (5 mW·cm$^{-2}$) for 30 minutes and then transferred to cells in 6 cm dishes. The dishes were divided into three groups. The first group was intermittently irradiated at 430 nm (1 mW·cm$^{-2}$; 10 seconds every 200 seconds) for 24 hours and then analyzed for cell cycle. The second group was irradiated for 24 hours and then transferred to the dark for 42 hours. The third group was maintained under light for 66 hours.

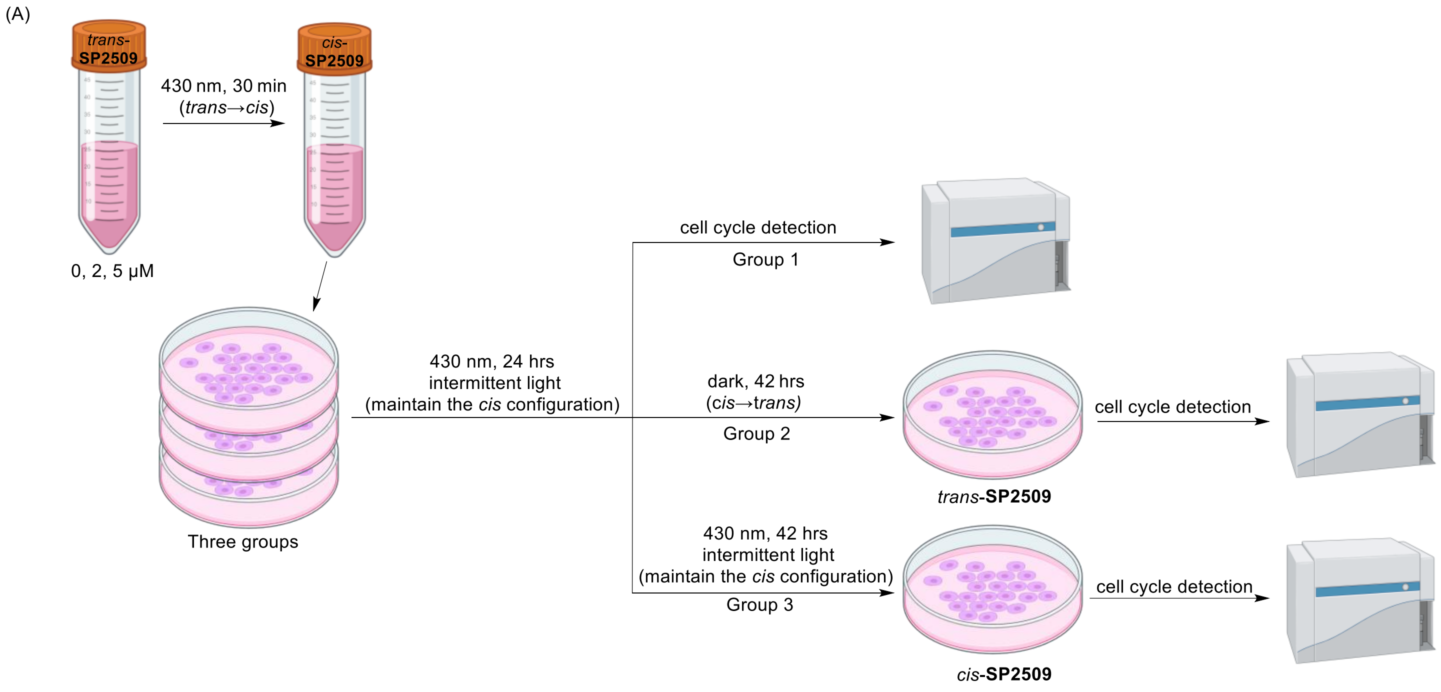


**Fig. S12.** (A) Schematic of the reciprocal *cis*-to-*trans* switching protocol and cell cycle detection. (B) Cell cycle in response to *cis*-**SP2509**, and *cis*-to-*trans* isomerization. The proportion of MGC-803 cells arrested in S phase remains comparable to control after 24 hours of *cis*-**SP2509** treatment, is markedly increased upon spontaneous *cis*-to-*trans* conversion in the dark for an additional 42 hours, and stays unchanged after 66 hours of continuous *cis*-**SP2509** exposure. (*to be continued*)

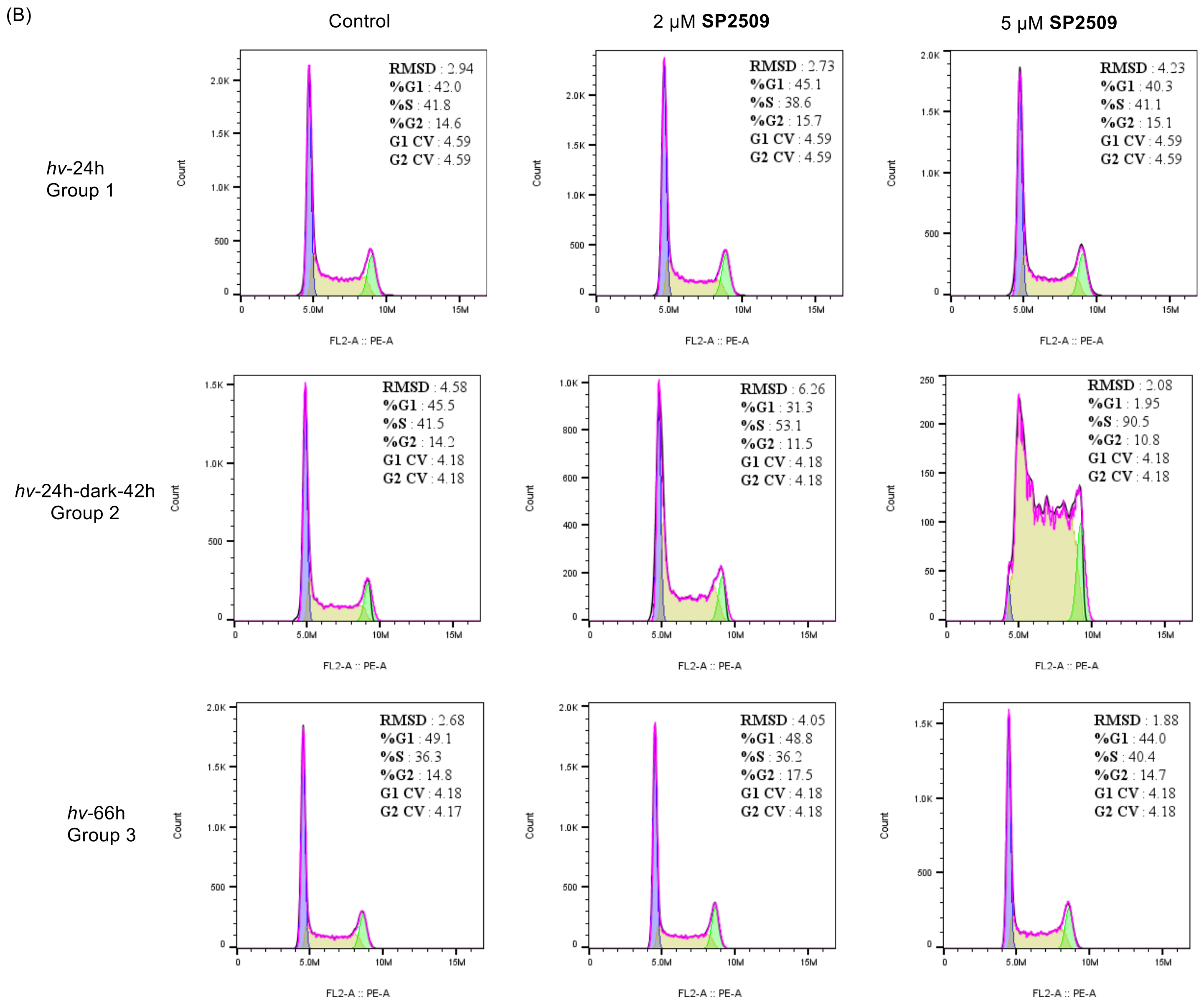


**Fig. S12.** (A) Schematic of the reciprocal *cis*-to-*trans* switching protocol and cell cycle detection. (B) Cell cycle in response to *cis*-**SP2509**, and *cis*-to-*trans* isomerization. The proportion of MGC-803 cells arrested in S phase remains comparable to control after 24 hours of *cis*-**SP2509** treatment, is markedly increased upon spontaneous *cis*-to-*trans* conversion in the dark for an additional 42 hours, and stays unchanged after 66 hours of continuous *cis*-**SP2509** exposure.

#### Quantitative Reverse Transcription PCR

OCI-AML3 cells were maintained in Iscove's modified Eagle medium (IMEM; Gibco, #C12440500BT) supplemented with 10% FBS (Gibco) and 1% penicillin/streptomycin (Gibco) at 37 °C under 5% $CO_2$. OCI-AML3 cells ($2 \times 10^6$) were seeded into each of four 10 cm cell culture dishes. After 12 hours, SP2509 at the indicated concentration (0 or 5 µM) was added to two dishes, which were then incubated in the dark for 24 h at 37 °C. For the light-treated groups, media containing *trans*-**SP2509** (0 or 5 µM) were pre-irradiated at 430 nm

(5 mW·cm$^{-2}$) for 30 minutes (to convert *trans*-**SP2509** to *cis*-**SP2509**) and then transferred to the remaining two dishes. These dishes were intermittently irradiated with 430 nm light (0.5 mW·cm$^{-2}$; 10 seconds every 200 seconds) and incubated for 24 hours at 37 °C. After the indicated treatments, total RNA was isolated and reverse transcribed. Quantitative real-time PCR analysis of BCL2A1, MYC, BIRC3 and CCND1 expression was performed using complementary DNA and TaqMan probes. Relative mRNA expression was normalized to GAPDH expression.

**Table S3.** Primers and TaqMan probes used in this study.

| **Gene** | **Forward sequence (5'-3')** | **Reverse sequence/probe (5'-3')** |
|---|---|---|
| GAPDH | TCGGAGTCAACGGATTTGGT | TTCCCGTTCTCAGCCTTGAC |
| | FAM-GGATATTGTTGCCATCAATGACCCC-BHQ1 | |
| BCL2A1 | ACCTAAATCTGGCTGGATGACT | GCCGGTTTCACAATATGGAGTG |
| | FAM-TGCTATCTCTCCTGAAGCAATACTGTTG-BHQ1 | |
| MYC | CTTCTCTCCGTCCTCGGATTCT | GAAGGTGATCCAGACTCTGACCTT |
| | FAM-CCGCAGGGCAGCCCCGAGC-BHQ1 | |
| BIRC3 | GCTTTTGCTGTGATGGTGGA | TGGCTTGAACTTGACGGATG |
| | FAM-CTGGAGATGATCCATGGGTTCAACAT-BHQ1 | |
| CCND1 | TGGAGCCCGTGAAAAAGAGC | TCTCCTTCATCTTAGAGGCCAC |
| | FAM-CCTGCAGCTGCTGGGGGC-BHQ1 | |

**Flow Cytometric Detection of Apoptosis**

OCI-AML3 cells were maintained in IMEM (Gibco, #C12440500BT) supplemented with 10% FBS (Gibco) and 1% penicillin/streptomycin (Gibco) at 37 °C under 5% $CO_2$. Cell apoptosis was detected using an Annexin V-FITC/PI apoptosis detection kit (Beyotime, #C1062). OCI-AML3 cells ($1.5 \times 10^6$) were seeded into each well of two 6-well plates. After 12 hours, *trans*-**SP2509** at the indicated concentrations (0, 0.5, 1, 2, 5 or 10 μM) was added to each well of one 6-well plate, and the plate was incubated in the dark for 48 hours at 37 °C. For the light-treated plate, media containing the corresponding concentrations of *trans*-**SP2509** were pre-irradiated at 430 nm (5 mW·cm$^{-2}$) for 30 minutes (to convert *trans*-**SP2509** to *cis*-**SP2509**) and then transferred to the second 6-well plate. The light-treated plate was intermittently irradiated with 430 nm light (0.5 mW·cm$^{-2}$; 10 seconds every 150 seconds) and incubated for 48 hours at 37 °C. After 48 hours treatment, apoptosis was measured with the Annexin V-FITC/PI detection kit. Briefly, cells were collected, washed with PBS, counted and resuspended in 195 μL binding buffer containing 5 μL Annexin V-FITC and 10 μL PI. The suspensions were incubated for 15 min at room temperature and analyzed on a FACSCalibur flow cytometer (BD). Data were analyzed using FlowJo software. FITC-positive events were defined as apoptotic cells, and the apoptotic fraction was calculated as the sum of the percentages of events in the Q2 and Q3 regions.

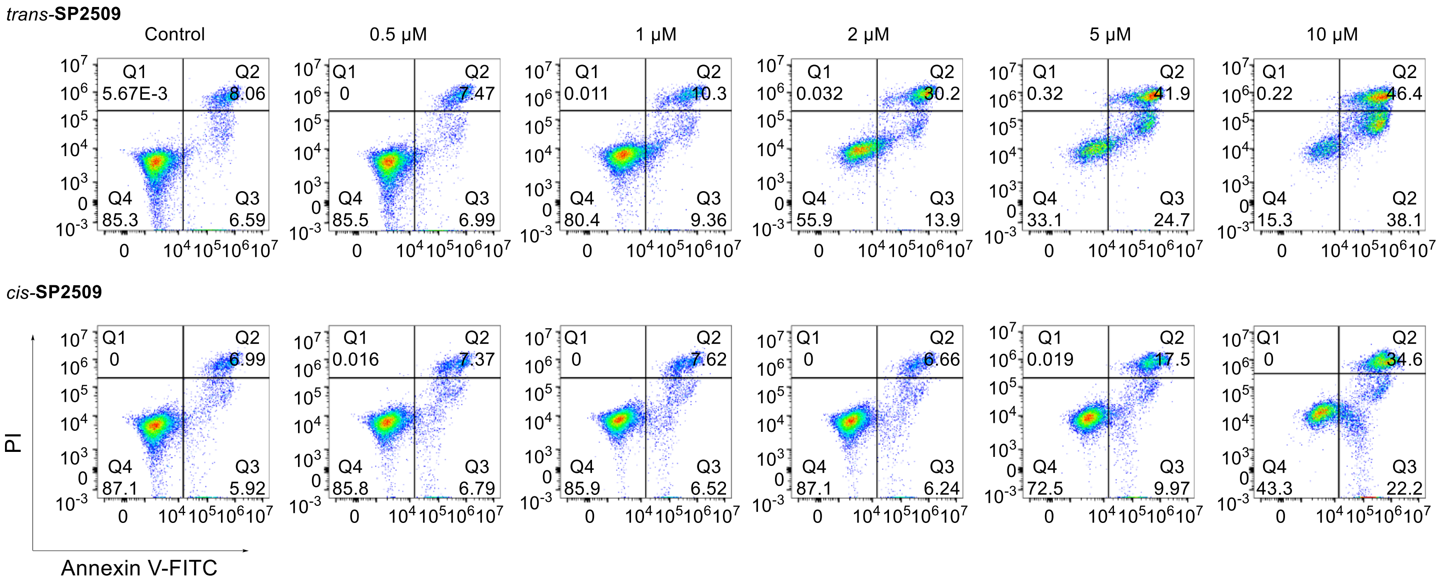


**Fig. S13.** Annexin V-FITC/PI flow-cytometry plots of OCI-AML3 cells treated with increasing concentrations of *trans*-**SP2509** or *cis*-enriched **SP2509** for 48 hours.

The experimental procedure for optically controlled dynamic regulation of apoptosis by **SP2509** is as follows: OCI-AML3 cells were seeded at a density of $1.5 \times 10^6$ cells per 6 cm dish, for a total of six dishes. The dishes were then divided into three treatment groups (two dishes each). In group 1, cells were treated with 0 or 5 μM *trans*-**SP2509** and cultured in the dark for 72 hours. In group 2, cells were treated with 0 or 5 μM *trans*-**SP2509** and cultured under constant light for 72 hours. For the light-treated plate, media containing the corresponding concentrations of *trans*-**SP2509** were pre-irradiated at 430 nm (5 $mW \cdot cm^{-2}$) for 30 minutes (to convert *trans*-**SP2509** to *cis*-**SP2509**) and then transferred to those dishes. The light-treated plate was intermittently irradiated with 430 nm light (0.5 $mW \cdot cm^{-2}$; 10 seconds every 150 seconds) and incubated at 37 °C. In group 3, both dishes received 5 μM *trans*-**SP2509** but were subjected to alternating light/dark cycles: one dish was incubated in the dark for 12 hours, then pre-irradiated at 430 nm (5 $mW \cdot cm^{-2}$) for 30 minutes before being exposed to light for 36 hours, and finally returned to the dark for 24 hours; the other dish was incubated under light for 24 hours, followed by dark for 24 hours, and then pre-irradiated at 430 nm (5 $mW \cdot cm^{-2}$) for 30 minutes before being exposed to light for 24 hours. At 12, 24, 36, 48, and 72 hours, cells in each dish were gently resuspended by pipetting, and aliquots of the cell suspension were collected for apoptosis analysis. Specifically, 0.3 mL was taken from dishes treated with 0 μM **SP2509**, and 0.5 mL was taken from those treated with 5 μM **SP2509**. Briefly, cells were collected, washed with PBS, counted and resuspended in 195 μL binding buffer containing 5 μL Annexin V-FITC and 10 μL PI. The suspensions were incubated for 15 minutes at room temperature and analyzed on a FACSCalibur flow cytometer (BD). Data were analyzed using FlowJo software. FITC-positive events were defined as apoptotic cells, and the apoptotic fraction was calculated as the sum of the percentages of events in the Q2 and Q3 regions.

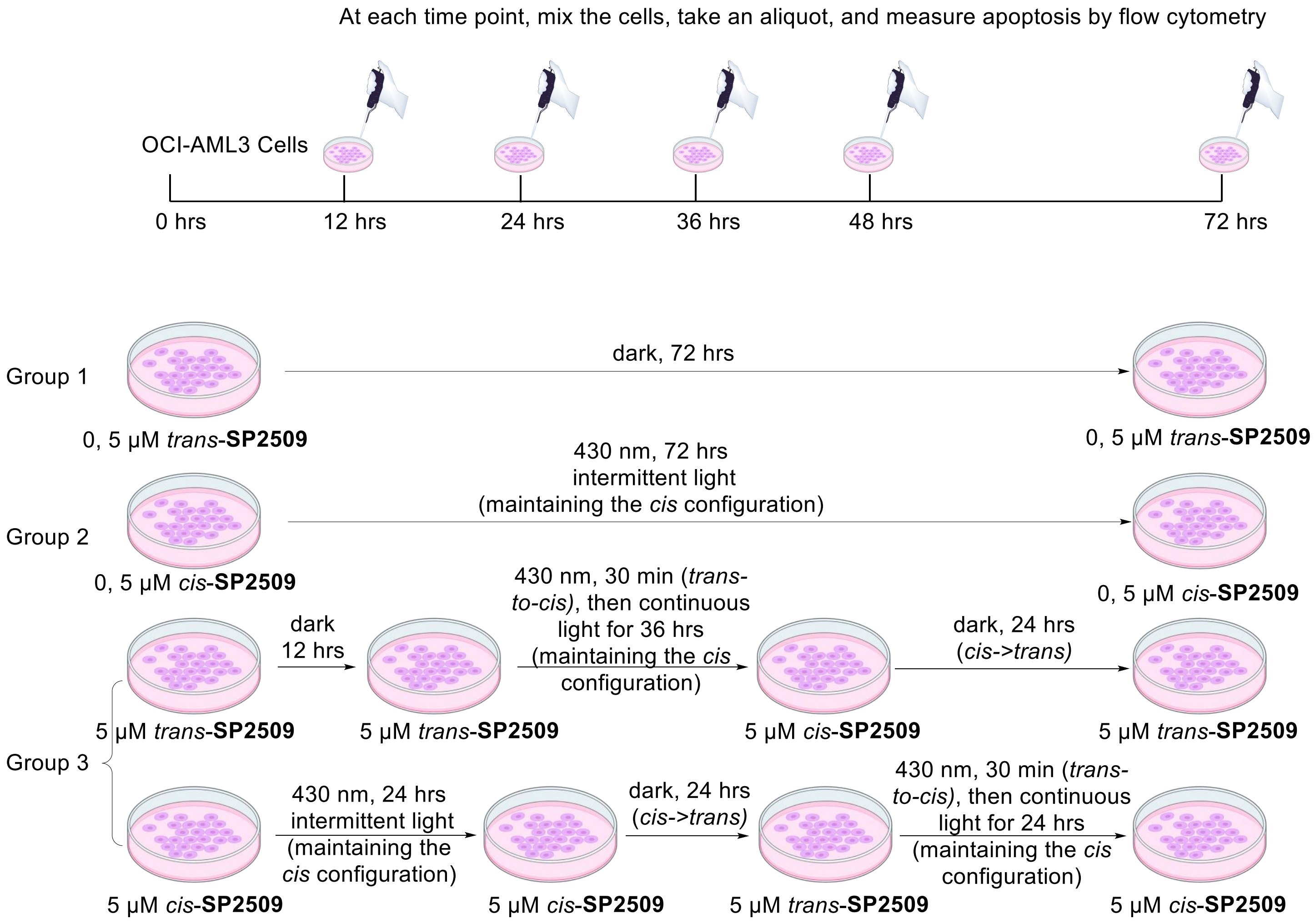


**Fig. S14.** Schematic of optically controlled dynamic regulation of apoptosis by **SP2509**.

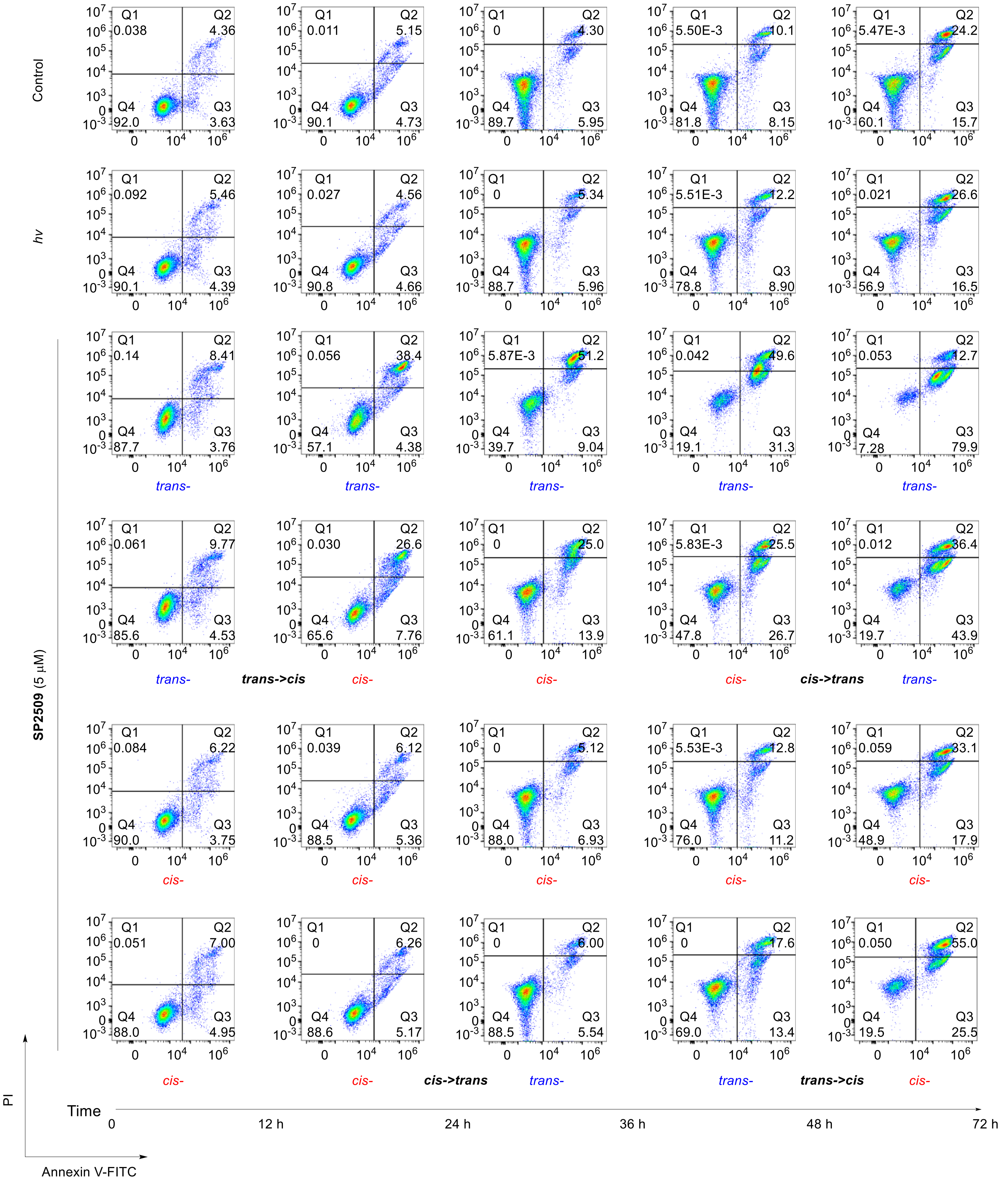


**Fig. S15.** Annexin V-FITC/PI flow cytometry plots showing that optical switching of **SP2509** dynamically regulates apoptotic commitment.

**RNA-seq Analysis**

OCI-AML3 cells were maintained in IMEM (Gibco, #C12440500BT) supplemented with 10% FBS (Gibco) and 1% penicillin/streptomycin (Gibco) at 37 °C under 5% $CO_2$. OCI-AML3 cells ($2 \times 10^6$) were seeded into each of four 10 cm cell culture dishes. After 12 hours, *trans*-**SP2509** at the indicated concentration (0 or 5 μM) was added to two dishes, which were then incubated in the dark for 24 hours at 37 °C. For the light-treated groups, media containing *trans*-**SP2509** (0 or 5 μM) were pre-irradiated at 430 nm (5 $mW \cdot cm^{-2}$) for 30 minutes (to convert it to *cis*-**SP2509**) and then transferred to the remaining two dishes. These dishes were intermittently irradiated with 430 nm light (0.5 $mW \cdot cm^{-2}$; 10 seconds every 200 seconds) and incubated for 24 hours at 37 °C. Three biological replicates of OCI-AML3 cells treated with 5 μM *trans*-**SP2509** or DMSO under dark or light conditions were prepared. Cells were washed three times with PBS, and cell pellets were collected, frozen in liquid nitrogen and sent to Beijing Novogene Co., Ltd. for RNA sequencing. RNA-seq libraries were prepared using the Fast RNA-seq Lib Prep Kit V2 (ABclonal, Cat. No. RK20306) and sequenced on an Illumina NovaSeq X Plus system to generate 150 bp paired-end reads. RNA-seq reads were aligned to the reference genome (ebi-39-homo-sapiens-grch38-primary-assembly) using HISAT2. Differential gene expression was analyzed using DESeq2. Genes with an absolute log2 fold change greater than 1 and padj ≤ 0.05 were considered differentially expressed. Gene Ontology (GO) and Kyoto Encyclopedia of Genes and Genomes (KEGG) pathway analyses were performed using clusterProfiler.

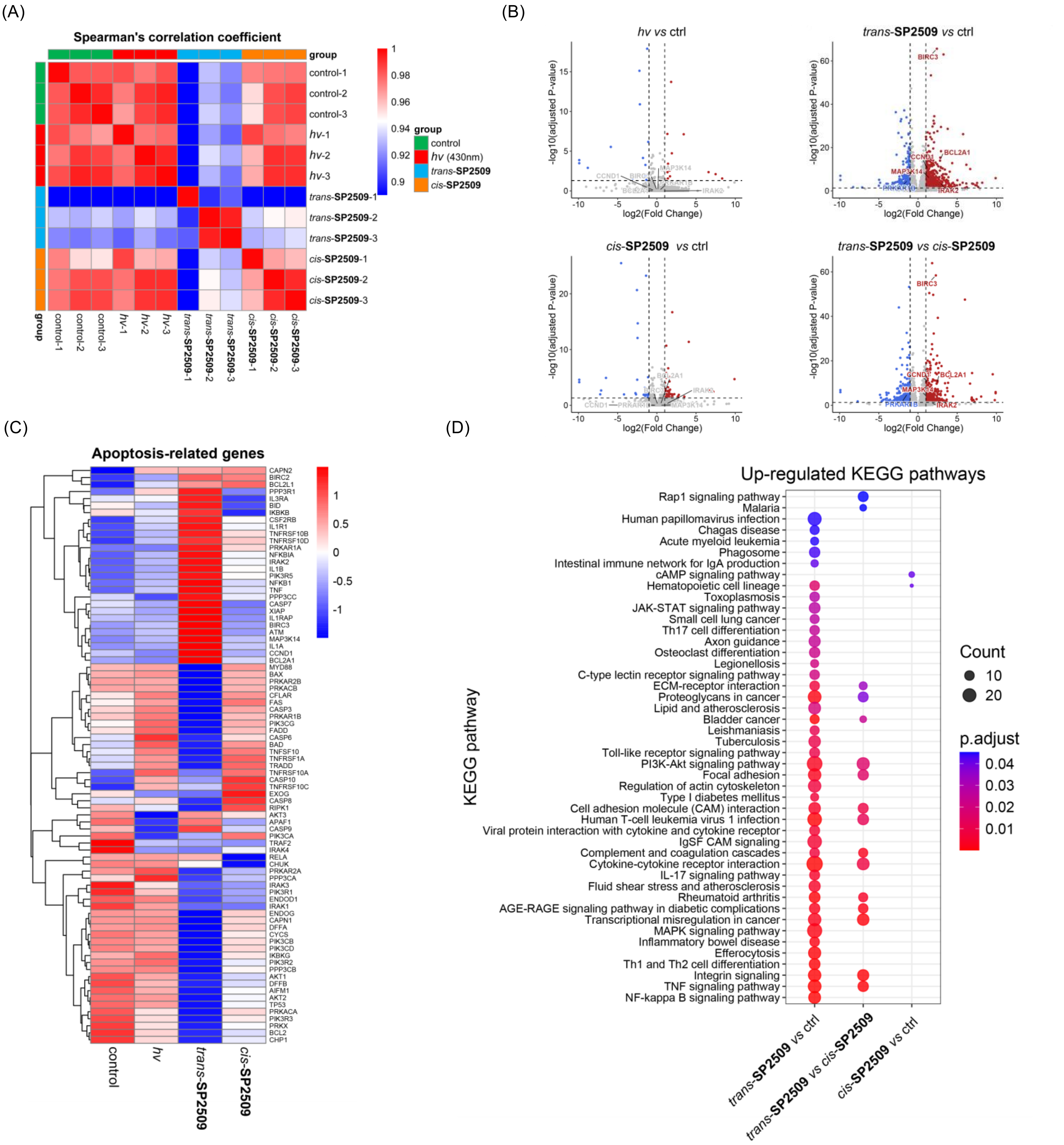


**Fig. S16.** (A) Sample-correlation analysis; (B) Volcano plot showing differentially expressed genes ($|\log_2$ fold change$| \geq 1$, adjusted P value $\leq 0.05$); (C) Heatmaps of apoptosis-related genes across group averages; (D) KEGG analysis of upregulated genes.

### NMR Spectrum

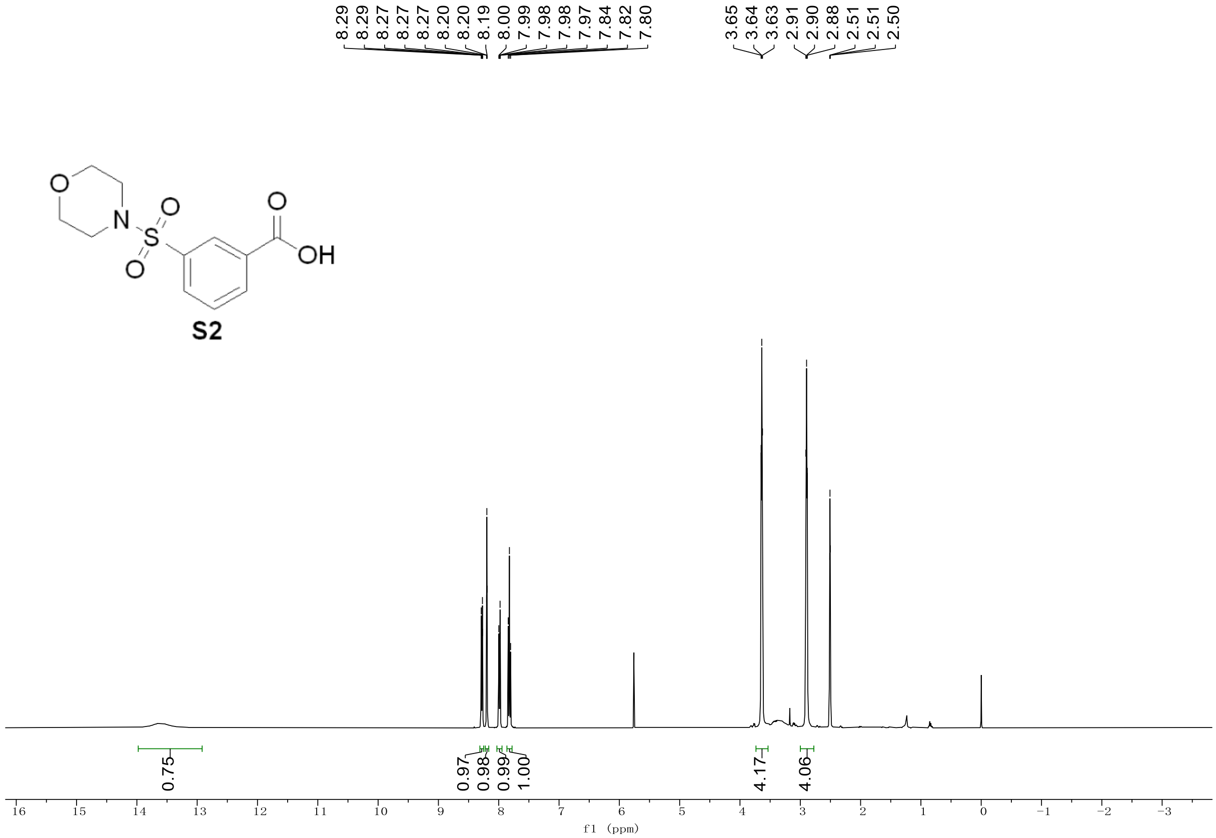


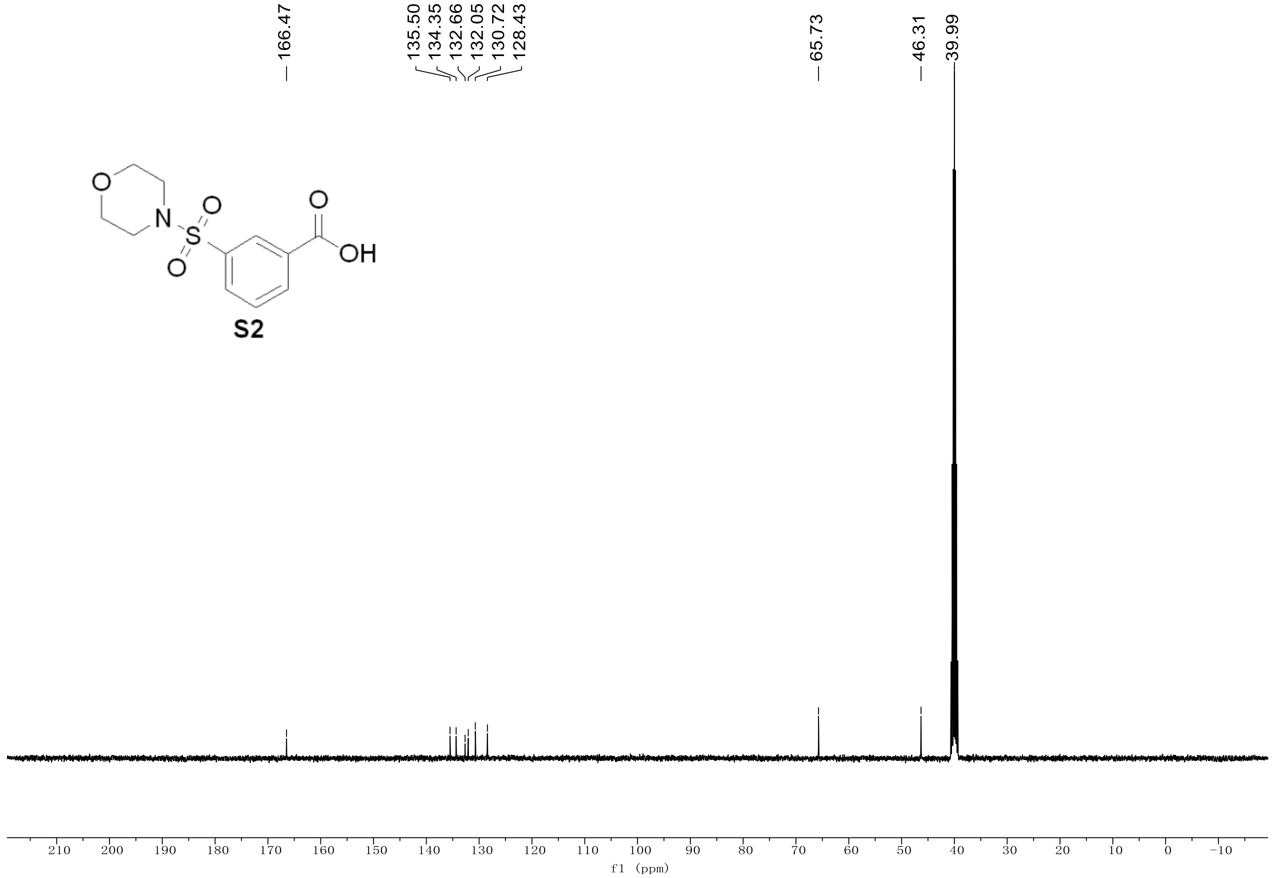

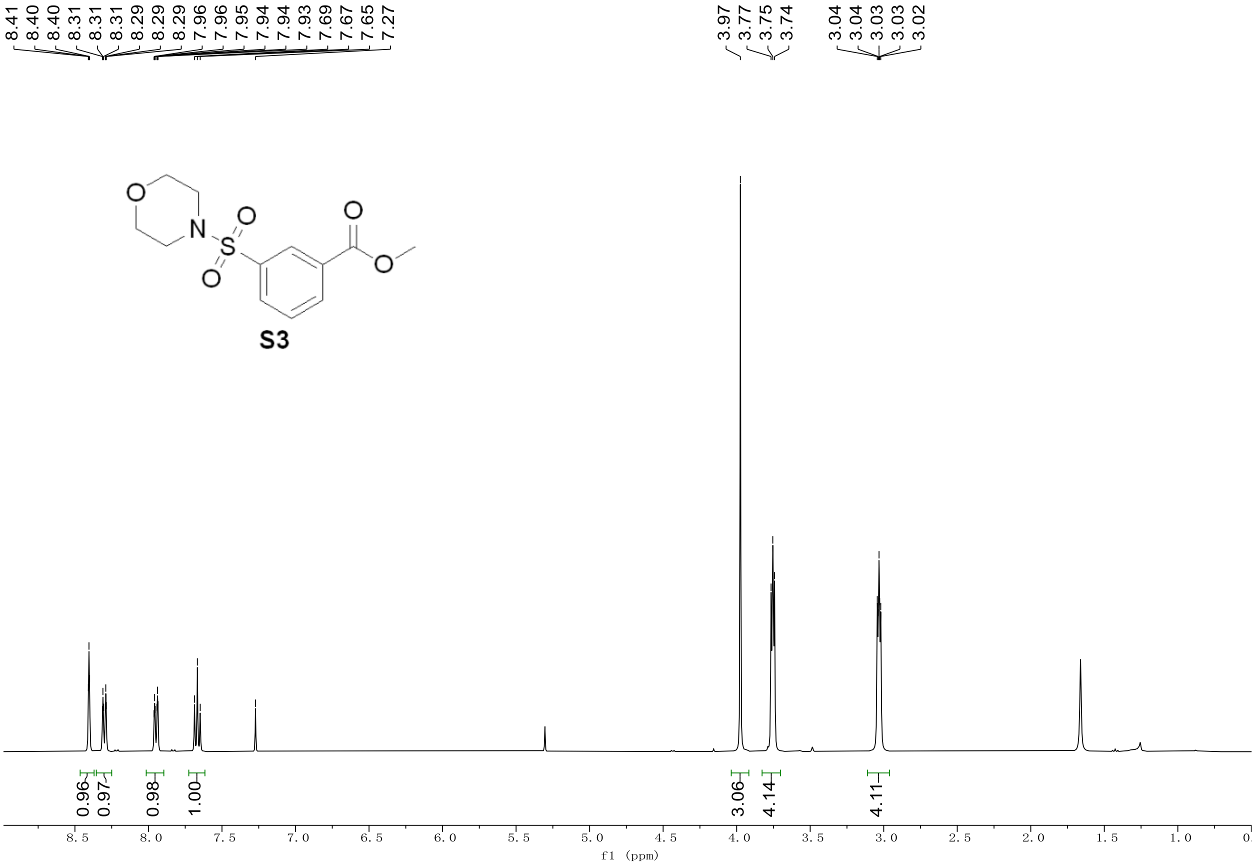
8.41
8.40
8.40
8.31
8.31
8.31
8.29
8.29
7.96
7.96
7.95
7.94
7.94
7.93
7.69
7.67
7.65
7.27
3.97
3.77
3.75
3.74
3.04
3.04
3.03
3.03
3.02
S3
0.96
0.97
0.98
1.00
3.06
4.14
4.11
f1 (ppm)

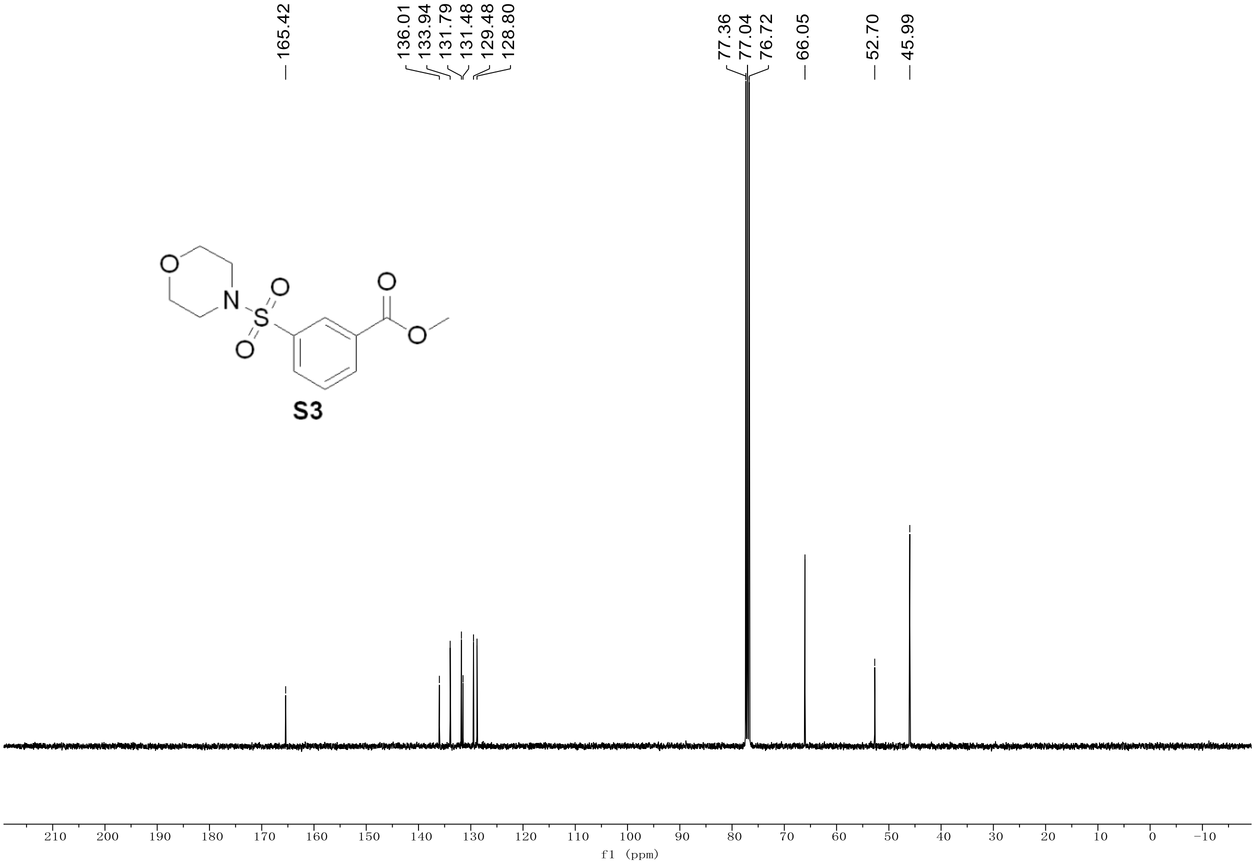
165.42
136.01
133.94
131.79
131.48
129.48
128.80
77.36
77.04
76.72
66.05
52.70
45.99
S3
f1 (ppm)

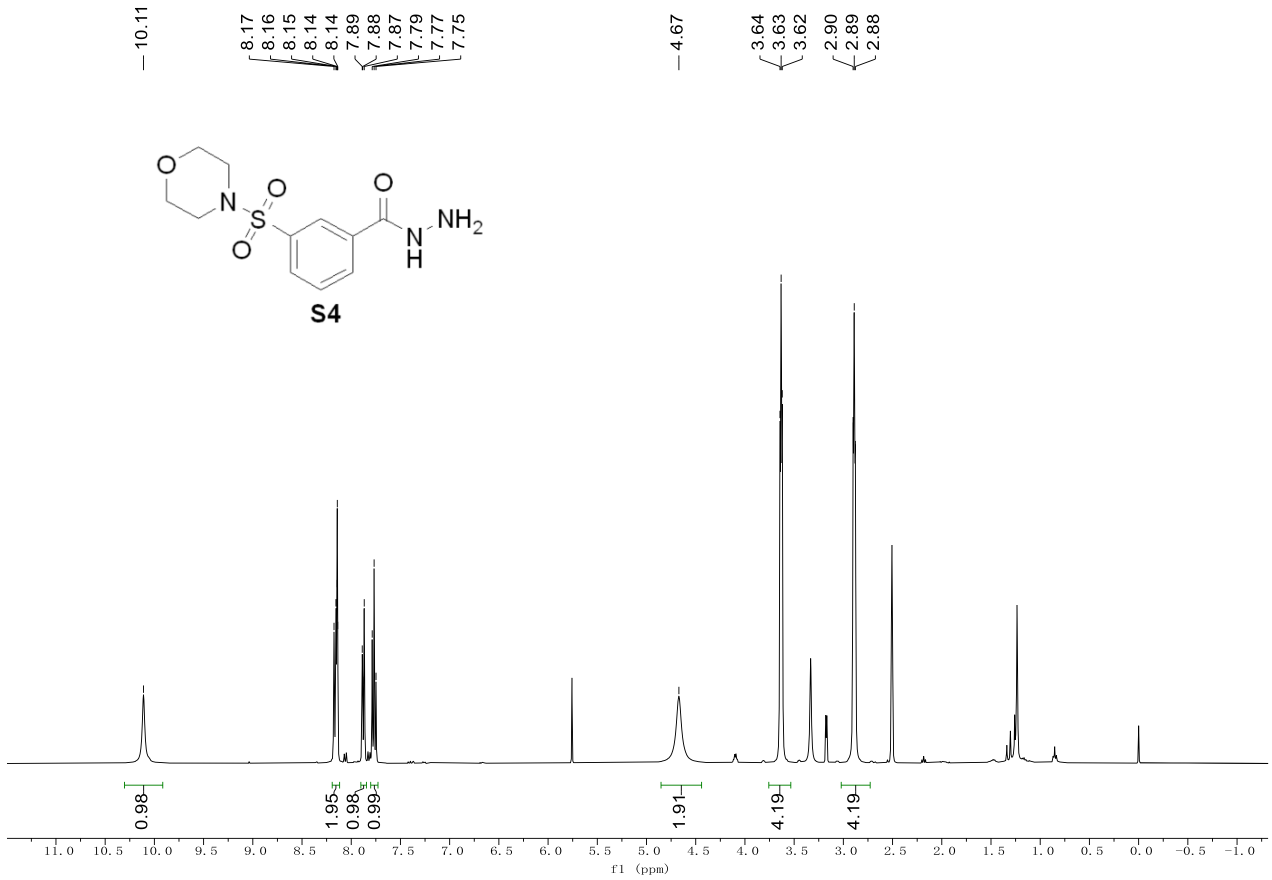

10.11
8.17
8.16
8.15
8.14
8.14
7.89
7.88
7.87
7.79
7.77
7.75
4.67
3.64
3.63
3.62
2.90
2.89
2.88
O
N
S
O
O
O
N
H
NH2
S4
0.98
1.95
0.98
0.99
1.91
4.19
4.19
f1 (ppm)


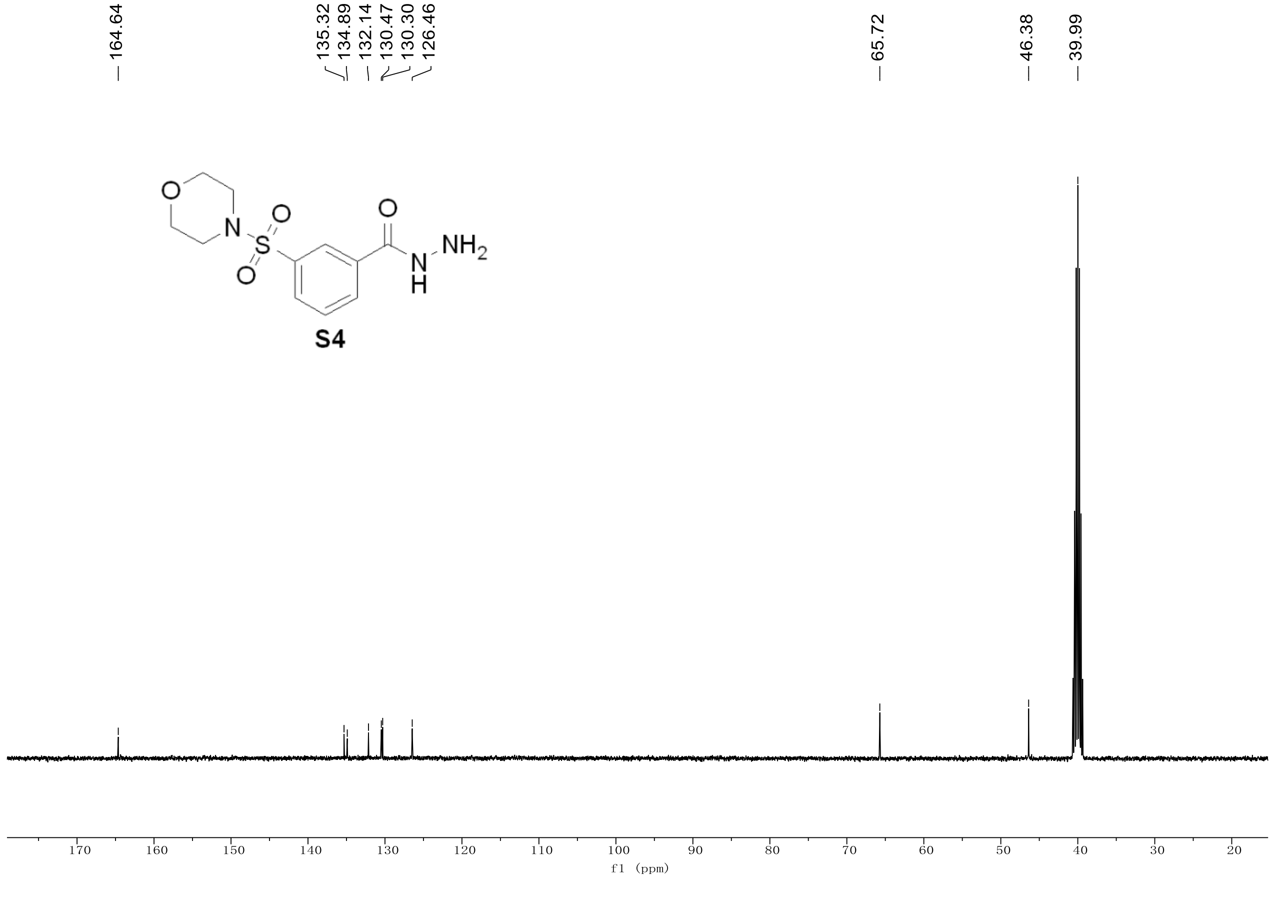

164.64
135.32
134.89
132.14
130.47
130.30
126.46
65.72
46.38
39.99
O
N
S
O
O
O
N
H
NH2
S4
f1 (ppm)

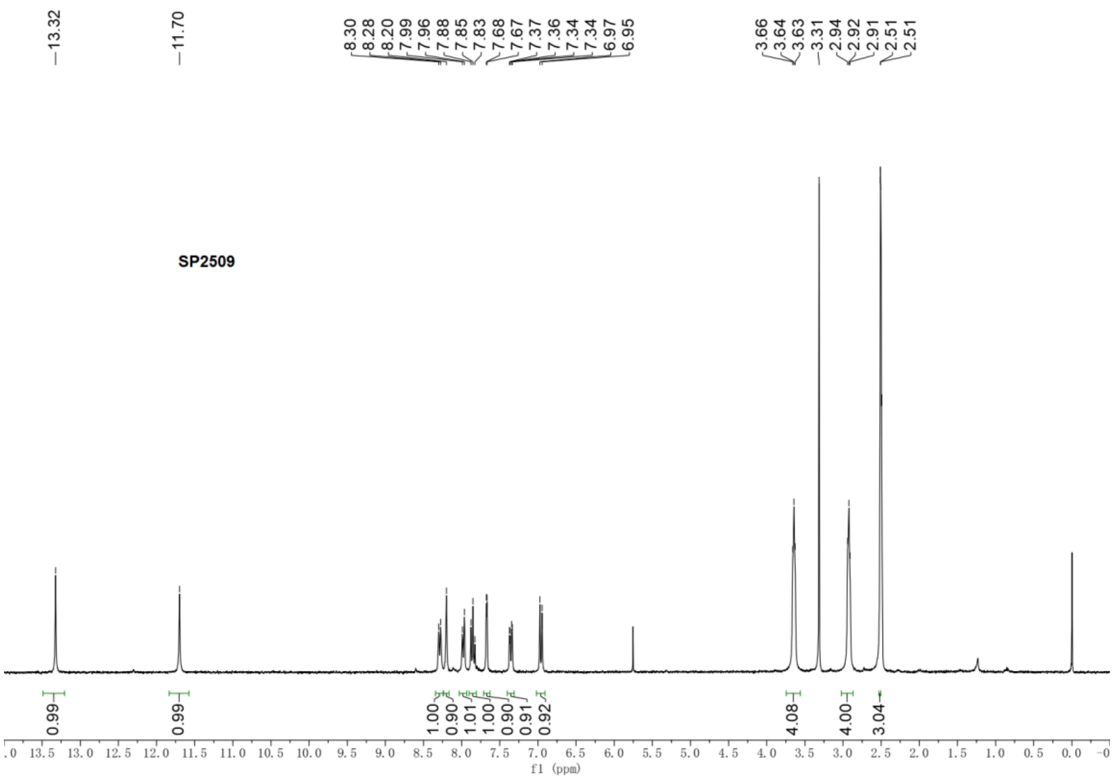
SP2509
f1 (ppm)
